# Alveolar epithelium propagates mechanical signals that control multi-cellular homeostasis in the lung

**DOI:** 10.1101/2025.06.04.657878

**Authors:** Chen Shen, Michael P. Morley, Sarah E. Schaefer, Gan Zhao, Joseph D. Planer, Ullas V. Chembazhi, Dakota L. Jones, Mijeong Kim, Yun Ying, Su Zhou, Shanru Li, Hannah Hallquist, Ana Lange, Maria C. Basil, Edward E. Morrisey

## Abstract

Respiratory motion imposes a constant mechanical strain that has important but poorly defined impact on tissue niches in the lung. We developed a reversible bronchial ligation model to induce and reverse unilateral blockade of lung mechanical motion *in vivo* and show that this leads to transcriptomic changes in multiple cell lineages that are not normalized upon reinitiation of respiratory motion. Perturbation of mechanosignaling specifically in alveolar epithelial type I (AT1) cells alters the transcriptomic state and fate of their niche neighbors, demonstrating that AT1 cells act as a node that propagates a mechanical cascade throughout the lung alveolus. Mechanically perturbed AT1 cells induce a distinct capillary endothelial cell state that persists after reactivation of respiratory motion, which is mediated by an integrin/TGF-β network within the alveolus that is vulnerable to pharmacological intervention. Importantly, AT1 mechanosignaling and intercellular communication are altered in chronic human lung diseases, highlighting the critical role of an AT1-driven mechanosensing network in lung disease biology. Thus, mechanosensing cells propagate biophysical signals that regulate tissue function and program tissue responses in disease.

## Introduction

Pulmonary gas exchange occurs within the alveoli, the elastic air sacs in the distal lung. During respiration, the alveoli are subjected to a constant, cyclical mechanical strain, which facilitates gas diffusion across a tight interface formed by the alveolar epithelium and the endothelium of the adjacent capillary plexus, with a fused basement membrane in between these two cells^1,2^. In lung diseases, this mechanical load is often disrupted, represented by damaged alveolar tissue architecture and compliance^3^. Despite the importance of pulmonary mechanical homeostasis, how these forces regulate cell behavior and fate within the gas exchange interface and the impact this has on tissue structure and function is poorly understood.

Recent studies have shown that mechanical forces are essential for maintaining alveolar epithelial cell fate and function^4–7^. Alveolar type II (AT2) epithelial progenitors require proper mechanical forces to differentiate into alveolar type I (AT1) cells, which facilitate alveolar gas exchange with the capillary endothelium^4–7^. AT1 cells exhibit high mechanosignaling activity and depend on mechanotransduction to maintain their identity, with loss-of-function mechanosignaling leading to their reprogramming into AT2 cells^4,8^. We have previously demonstrated that AT1 cells are responsive to respiration-mediated mechanical forces *in vivo* by surgical blockade of lung motion^4^. Perturbed respiratory motion caused AT1 cells to reprogram into AT2 cells, mimicking the phenotype observed upon attenuation of mechanotransduction in AT1 cells through loss of Hippo signaling and actin dynamics^4,8^. However, in all of these models, loss of mechanical signaling was permanent, which did not allow for the characterization of the impact that reinitiation of this signaling process would have on cell fate and tissue dynamics.

Here, we developed a reversible bronchial ligation model to induce temporary unilateral blockade of lung motion in mice, followed by restoration of respiratory movements, and used this model to characterize both transient and persistent alterations in cell fate and behavior induced by respiration-mediated mechanical forces. This model has allowed us to generate a tissue-wide mechanotransduction map in the lung at single-cell resolution. Analyses of our single-cell RNA sequencing (scRNAseq) dataset demonstrate that temporary respiratory blockade can lead to persistent cell transcriptomic alterations, which can occur through mechanisms beyond intrinsic mechanotransduction in cells. These analyses reveal that AT1 cells are a node for mechanotransduction in the lung alveolus that senses biophysical responses and propagates mechanical signals to other niche neighbors including AT2 epithelial, alveolar fibroblast, and CAP1 endothelial progenitors. Deregulation of mechanotransduction specifically in AT1 cells induces an irreversible, transcriptionally distinct CAP state via intercellular integrin/TGF-β signaling originated from AT1s. Human lung scRNAseq data indicate aberrant AT1 mechanosensing in chronic lung diseases, including opposing effects on chronic obstructive pulmonary disease (COPD) and idiopathic pulmonary fibrosis (IPF). Together, our findings demonstrate a pivotal role for an AT1 mechanosensing network that propagates a wave of signaling throughout the alveolus critical for maintaining tissue integrity and can be used to phenotype human lung diseases.

## Results

### Mechanical forces maintain multi-cellular homeostasis of the lung alveolus

To understand how cells in the lung respond to changes in mechanical forces, we developed a reversible bronchial ligation model to induce and reverse blockade in respiration-induced mechanical forces *in vivo* (**Fig. 1a**)^4^. In this model, the left main bronchus of mice is ligated by a micro-clip to temporarily restrain unilateral respiratory movements (Ligation, **Fig. 1a**), which can be later removed to restore respiratory motion (Reversed, **Fig. 1a**). Reversed lungs were found to be reinflated and their histological appearance by Hematoxylin and Eosin staining was relatively normal (**Extended Data Fig. 1a-b**). We performed sham, ligation, or reversed surgery in adult wildtype mice, isolated cells from the lungs and performed single-cell RNA sequencing (scRNA seq) to determine transcriptomic changes in cells due to modulated mechanical forces (**Fig. 1b**). All expected cell types were successfully captured in the scRNAseq experiment and annotated (**Fig. 1c-f and Extended Data Fig. 1c**). We noticed altered transcriptome in various resident cell types within the alveolus in response to restrained respiration, especially in progenitor cells such as AT2 epithelial cells, CAP1 capillary endothelial cells and alveolar fibroblasts (**Fig. 1c-f**). Of note, not all transcriptomic changes in these cell types were fully normalized after restored respiratory motion. This suggests that disrupted mechanical forces can induce irreversible transcriptomic changes in the lung even after normalization of respiration, with a potential impact on progenitor cell function.

**Figure 1:**
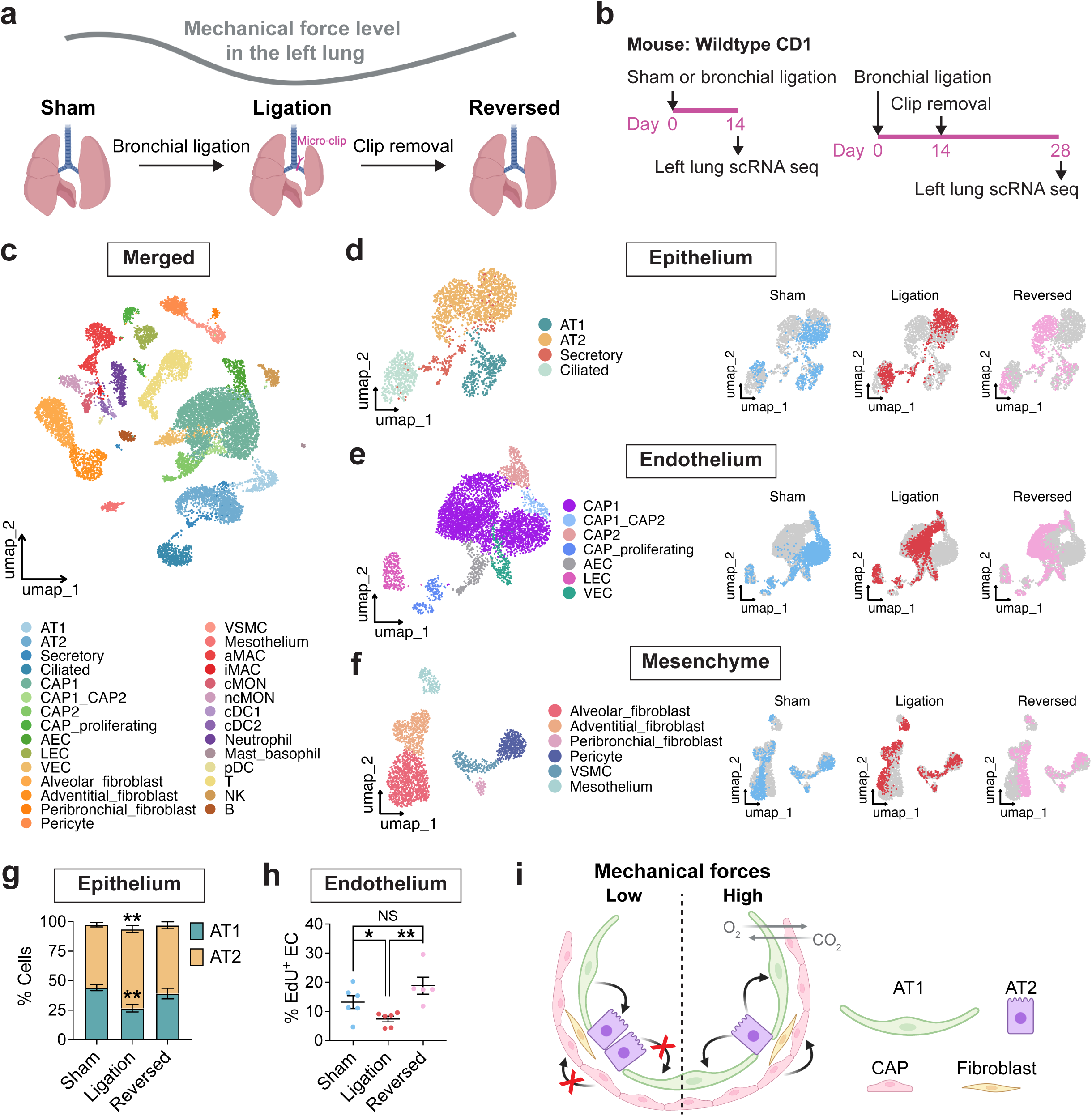
Respiration-mediated mechanical forces maintain multi-cellular homeostasis of the lung alveolus. **a.** A schematic of reversible bronchial ligation model that modulates unilateral lung motion. **b.** A schematic of scRNA seq experiment. Cells from wildtype CD1 mouse lungs after sham, bronchial ligation or reversed surgery were isolated and used for scRNA seq analyses. **c.** A merged UMAP plot of distal lung cells from sham, ligated and reversed mouse lungs. Cells were annotated by cell type. **d-f.** UMAP plots of cells from indicated lineages annotated by cell type and treatment. **g.** Percentage of AT1 and AT2 cells in indicated mouse lungs (n = 4 or 5, mean ± S.E.M.). Asterisks indicate statistical significance (** *p* value < 0.01, unpaired two-tailed Student’s *t* test). **h.** Percentage of EdU incorporated endothelial cells in indicated mouse lungs (n = 5 or 6, mean ± S.E.M.). Asterisks indicate statistical significance (* *p* value < 0.05, ** *p* value < 0.01, unpaired two-tailed Student’s *t* test). NS: no statistical significance. **i.** A model depicting mechanical force-induced cell responses in the lung alveolus.

The facultative progenitor AT2 cells self-renew and differentiate into AT1 epithelial cells upon lung injury^9–13^. Prior studies have demonstrated that mechanical forces are critical for AT2-to-AT1 differentiation^4–7^. In addition, AT1 cells are sensitive to mechanical strain and can reprogram into AT2 cells in response to inhibited mechanical signaling^4,8^. Immunostaining of mouse lung tissues using antibodies against AT1- or AT2-specific marker showed that compared to the sham procedure, ligation induced a significant decrease in AT1 cell proportion, while AT2 cells showed an increased percentage (**Fig. 1g and Extended Data Fig. 2a**). This is similar to what we have previously shown and in line with AT1-to-AT2 reprogramming upon blocked mechanical forces. After the procedure is reversed, AT1 and AT2 cell percentages were normalized, likely through AT2-AT1 differentiation (**Fig. 1g**). Gene set enrichment analysis showed that Wnt and FGFR signaling that are essential for AT2 progenitor cell function were downregulated in AT2 cells in ligated mouse lungs but restored after reversal (**Extended Data Fig. 2b**)^12,14^.

The lung capillary plexus and alveolar epithelium form a tight interface to facilitate efficient gas exchange^1,2^. Capillary endothelium (CAPs) is comprised of two distinct populations, CAP1 which are the proliferative progenitor subpopulation and CAP2 which exhibit a different phenotype consisting of increased expression of angiogenic genes^15,16^. The regeneration of CAPs has been shown to play a critical role in rebuilding lung tissue structure and function after acute injury and is driven largely by proliferation and differentiation of CAP1s^17–20^. To understand how progenitor CAPs respond to mechanical insults, we examined overall CAP proliferation by detecting cellular 5-ethynyl-2’-deoxyuridine (EdU) incorporation in mouse ligated lungs and those after reversed surgery (**Extended Data Fig. 3a-b**). We found that baseline EdU incorporation in CAPs was significantly reduced in the ligated mouse lungs (**Fig. 1h**). However, this was fully rescued when the ligation is reversed (**Fig. 1h**). Some CAP transcriptomic changes induced by ligation were reversed by ligation removal, such as changes in TGF-β signaling (**Extended Data Fig. 3c**). Together, these findings suggests that mechanical forces are essential for maintaining the multi-cellular integrity and function of the lung alveolus (**Fig. 1i**).

### Cellular response to mechanical forces can be independent of intrinsic mechanosignaling

One of the primary transducers of mechanical forces into cells is the Hippo pathway, which is controlled by two essential transcription co-factors, Yes-associated protein (YAP) and transcriptional coactivator with PDZ-binding motif (TAZ) (**Fig. 2a**)^21–23^. To evaluate the changes in cellular mechanosignaling, a transcriptional score was generated for each cell type based on the expression profile of 22 known YAP/TAZ target genes (mechanoscore) (**Fig. 2a**)^24^. In the homeostatic lung, cells exhibit differential levels of mechanotransduction based on this score, with AT1 cells and mesenchymal cells showing the highest mechanosignaling activity relative to other cell types (mech-high cells, **Fig. 2b-d**). Using Euclidian distance analysis, we quantified transcriptomic changes in lung cells after bronchial ligation and upon reversal, based on their proximity to baseline gene expression profiles. Ligation-induced transcriptomic changes were noted in various resident cell types within the alveoli, including AT1 cells and all mesenchymal cells, consistent with their sensitivity to this perturbation (**Fig. 2e-f and Extended Data Fig. 4a-b**)^4,25,26^. We noted that transcriptional alterations in mesenchyme cells persisted after reversed surgery, whereas the AT1 cell transcriptome was largely normalized, suggesting AT1 cells are highly adaptable to respiration-mediated mechanical forces (**Fig. 2e and Extended Data Fig. 4b**). Interestingly, some cells with low baseline mechanical activity (mech-low cells) also exhibited an altered transcriptomic profile in response to perturbed mechanical forces, e.g. AT2 cells and CAPs (**Fig. 2g-h and Extended Data Fig. 4a-b**). These changes were not distinct to mechanotransduction, suggesting that biophysical forces can regulate gene expression and cell state in these mech-low cells via mechanisms beyond classical mechanical signaling (**Fig. 2g-h and Extended Data Fig. 4a-b**). Together, these studies reveal how biophysical forces are sensed at both cell-intrinsic and -extrinsic levels to maintain tissue homeostasis in a mechanically loaded organ such as the lung.

**Figure 2:**
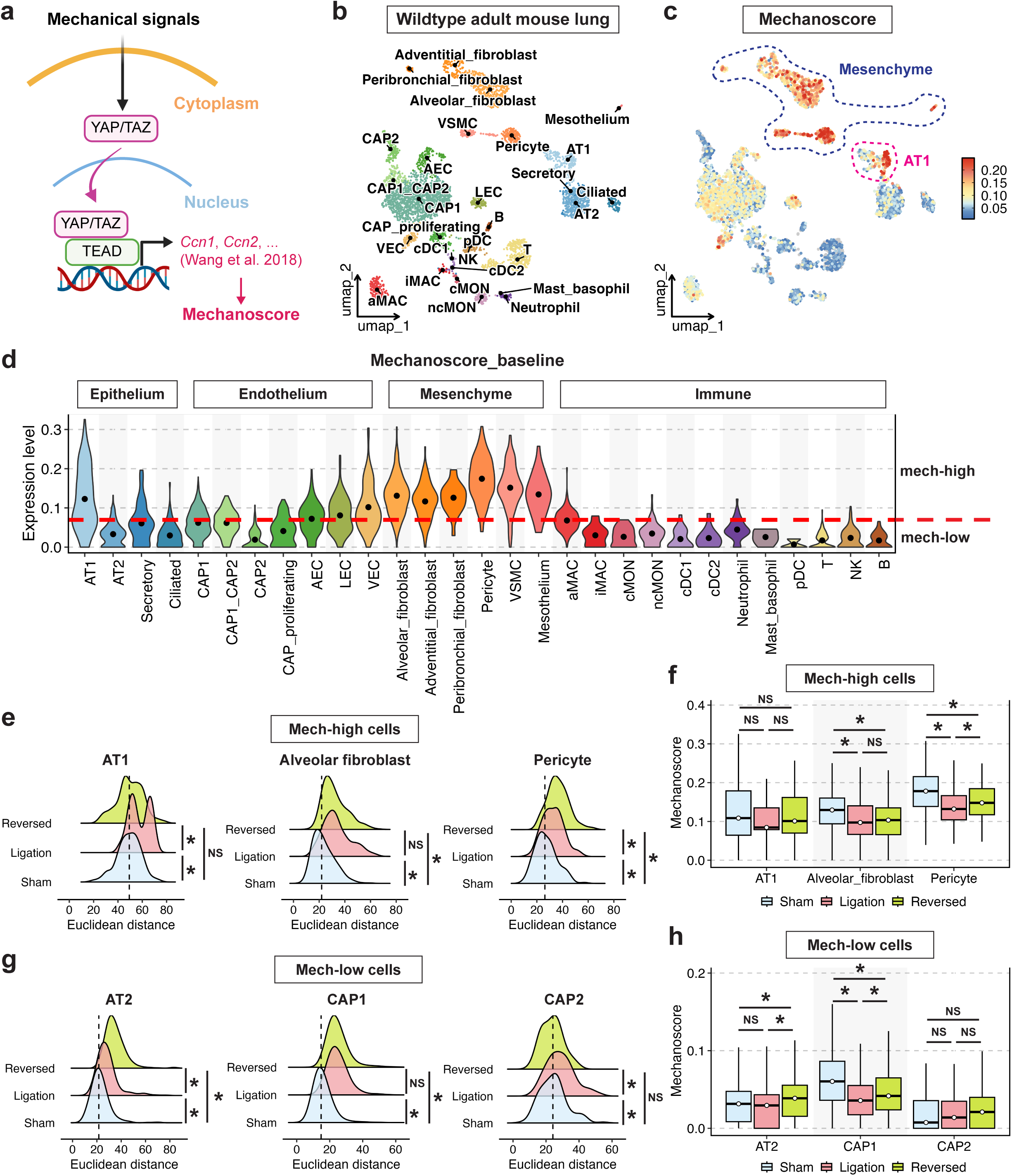
Alveolar cell responses to mechanical forces can be independent of intrinsic mechanosignaling. **a.** A schematic of the YAP/TAZ mechanosignaling pathway. 22 YAP/TAZ target genes reported in Wang et al. 2018 were used to generate cell mechanoscore. **b.** A UMAP plot of distal lung cells from sham mouse lungs. Cells were annotated by cell type. **c-d.** Baseline mechanoscore of each cell type in the distal lung. Red dashed line in (**d**) indicates the mean of mechanoscore across all cells. **e,g.** Euclidian distance indicating cellular transcriptional similarity across treatments. An asterisk indicates statistical significance (* *p* value < 0.05, pairwise Wilcoxon test). NS: no statistical significance. **f,h.** Mechanoscore of indicated cell types in sham, ligation, or reversed mouse lungs. An asterisk indicates statistical significance (* *p* value < 0.05, pairwise Wilcoxon test). NS: no statistical significance.

### AT1 cells are a node that propagates mechanical signaling cascades to neighboring cells to control niche homeostasis

Within the alveoli, the large squamous AT1 cells occupy over 90% of the surface area^1,2^. AT1 cells are adjacent to many other cells in the alveolar niche, including AT2 cells and CAPs, which play an essential role in lung structure and function and communicate with AT1 cells^1,2,27,28^. Loss of mechanical signaling in AT1 cells, either by loss of the pivotal cytoskeleton assembly gene *Cdc42* or through bronchial ligation, causes AT1 cells to reprogram into AT2 cells^4^. Therefore, we genetically deleted *Cdc42* in AT1 cells (*Cdc42*^AT1-KO^) to determine the impact from loss of mechanosignaling in AT1 cells on their alveolar neighbor cells using scRNAseq (**Fig. 3a**). Whole lung scRNAseq analysis revealed that loss of mechanotransduction in AT1 cells led to altered transcriptome of specific neighboring cell types, including AT2 cells, CAP1s and alveolar fibroblasts (**Fig. 3b-e and Extended Data Fig. 5a-b**), supporting the concept that AT1 cells are a node of mechanotransduction in the lung.

**Figure 3:**
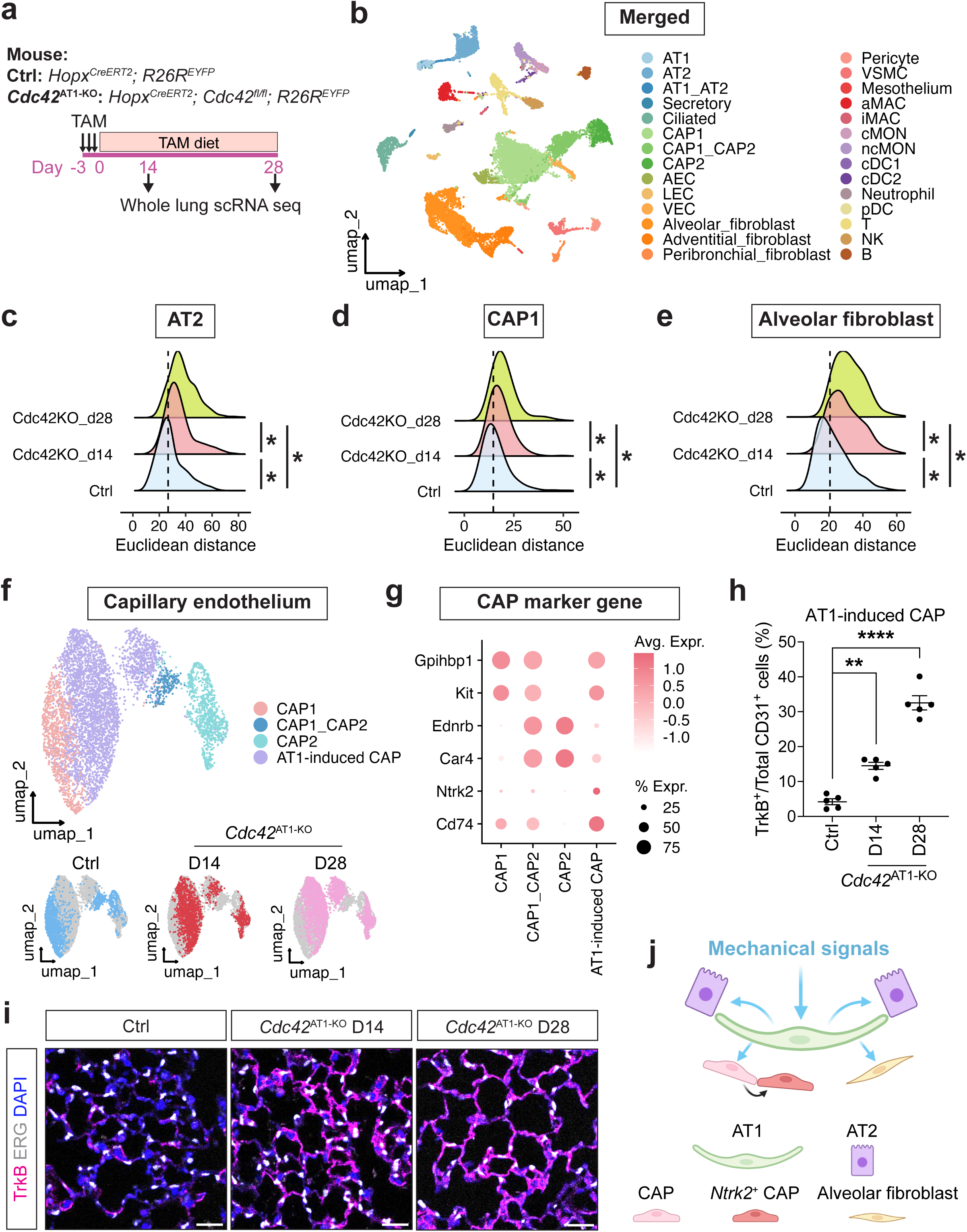
AT1 cells are a mechanotransduction node in the lung. **a.** A schematic of genetic attenuation of mechanotransduction in AT1s. Control AT1-trace mice or those with a *Cdc42^fl/fl^* allele were administrated with three doses of tamoxifen, followed by tamoxifen diet. Cells from control or *Cdc42*^AT1-KO^ mouse lungs were isolated and used for scRNA seq analyses at indicated time points. **b.** A merged UMAP plot of distal lung cells from control or *Cdc42*^AT1-KO^ mouse lungs. Cells were annotated by cell type. **c-e.** Euclidian distance indicating cellular transcriptional similarity across treatments. Asterisks indicate statistical significance (* *p* value < 0.05, pairwise Wilcoxon test). NS: no statistical significance. **f.** UMAP plots of CAPs from control and *Cdc42*^AT1-KO^ mouse lungs. Cells were annotated by cell type or treatment. **g.** A dot plot showing expression of indicated genes in each cell type. **h.** Flow cytometry results showing the percentage of AT1-induced CAP in *Cdc42*^AT1-KO^ mouse lungs after three doses of tamoxifen for 14 or 28 days, or control mouse lungs (n = 5, mean ± S.E.M.). Asterisks indicate statistical significance (** *p* value < 0.01, **** *p* value < 0.0001, unpaired two-tailed Student’s *t* test). **i.** Representative IHC images of control or *Cdc42*^AT1-KO^ mouse lungs stained with TrkB and ERG (scale bar = 25 μm). **j.** A model depicting AT1 cells as a node of mechanotransduction in the alveolus.

To further understand the impact of AT1 mechanoresponses on adjacent CAPs, we subseted CAPs from *Cdc42*^AT1-KO^ scRNAseq data to analyze their transcriptomic profile at a higher resolution (**Fig. 3b and f**). We identified a CAP state specific to *Cdc42*^AT1-KO^ lungs (AT1-induced CAP) that was characterized by enriched expression of a set of genes including *Ntrk2* and *Cd74* (**Fig. 3f-g**). To validate the presence of these cells, we isolated cells from control or *Cdc42*^AT1-KO^ mouse lungs and performed flow cytometry analyses using a Tropomyosin receptor kinase B (TrkB) antibody to quantitate the percentage of *Ntrk2^+^* CAPs. We confirmed that loss of *Cdc42* in AT1 cells led to the emergence of *Ntrk2^+^* CAPs in mouse lungs in a time-dependent manner (**Fig. 3h and Extended Data Fig. 6a**). In addition, tissue immunostaining using TrkB antibody validated the presence of *Ntrk2^+^* AT1-induced CAPs in mouse ligated lungs (**Fig. 3i**). These results indicate that AT1 cells can transmit their mechanosensitive responses to neighboring cells to mediate lung tissue homeostasis (**Fig. 3j**).

### The AT1 mechanosensing-induced CAP state is permanent and functionally distinct

Given the alteration in CAP gene expression upon loss of *Cdc42* in AT1 cells, we next asked whether this also occurred upon blockade of respiratory motion. We reclustered the CAP populations in the scRNAseq data from sham, ligated, or reversed mouse lungs to identify alterations in cell states (**Fig. 1b-c and Fig. 4a**). CAP1 progenitors and the intermediate CAP1_CAP2 cell state were significantly altered at the transcriptional level, losing their homeostatic state upon respiratory blockade (**Fig. 4a-b**). Instead, we identified a distinct population of CAPs that emerged in ligated mouse lungs and persisted after reversal (Ligation-induced CAP), with enriched expression of a number of genes induced by inhibiting mechanotransduction in AT1 cells, e.g. *Ntrk2* (**Fig. 4a-c, Extended Data Fig. 7a-b**). Moreover, we found that there was a strong correlation between the transcriptome of Ligation-induced CAPs and AT1-induced CAPs, suggesting that they represent similar cell states (**Fig. 4d**). We hereafter termed these *Ntrk2^+^*CAPs as mechanically induced CAPs (miCAPs). The existence of miCAPs was further validated in ligated and reversed mouse lungs by flow cytometry and tissue immunostaining using TrkB antibody (**Fig. 4e-f**). Despite the persistence of miCAP state in the reversed lung, we noted that these cells exhibited enhanced proliferation compared to TrkB^-^endothelial cells (ECs) during reversal of bronchial ligation (**Extended Data Fig. 3a and 8a-c**). To determine the fate change of miCAPs in this process, we used *Ntrk2^+^* lineage tracing to track miCAPs and noted that approximately 50% of the traced cells lost TrkB expression after restoration of respiratory motion (**Fig. 4g and Extended Data Fig. 8d-i**). Interestingly, a similar percentage of miCAPs were capable of differentiating into CAP2 cells upon ligation reversal (**Fig. 4h-i**). These results suggest that some miCAPs can differentiate into CAP2 cells during resolution of lung motion blockade.

**Figure 4:**
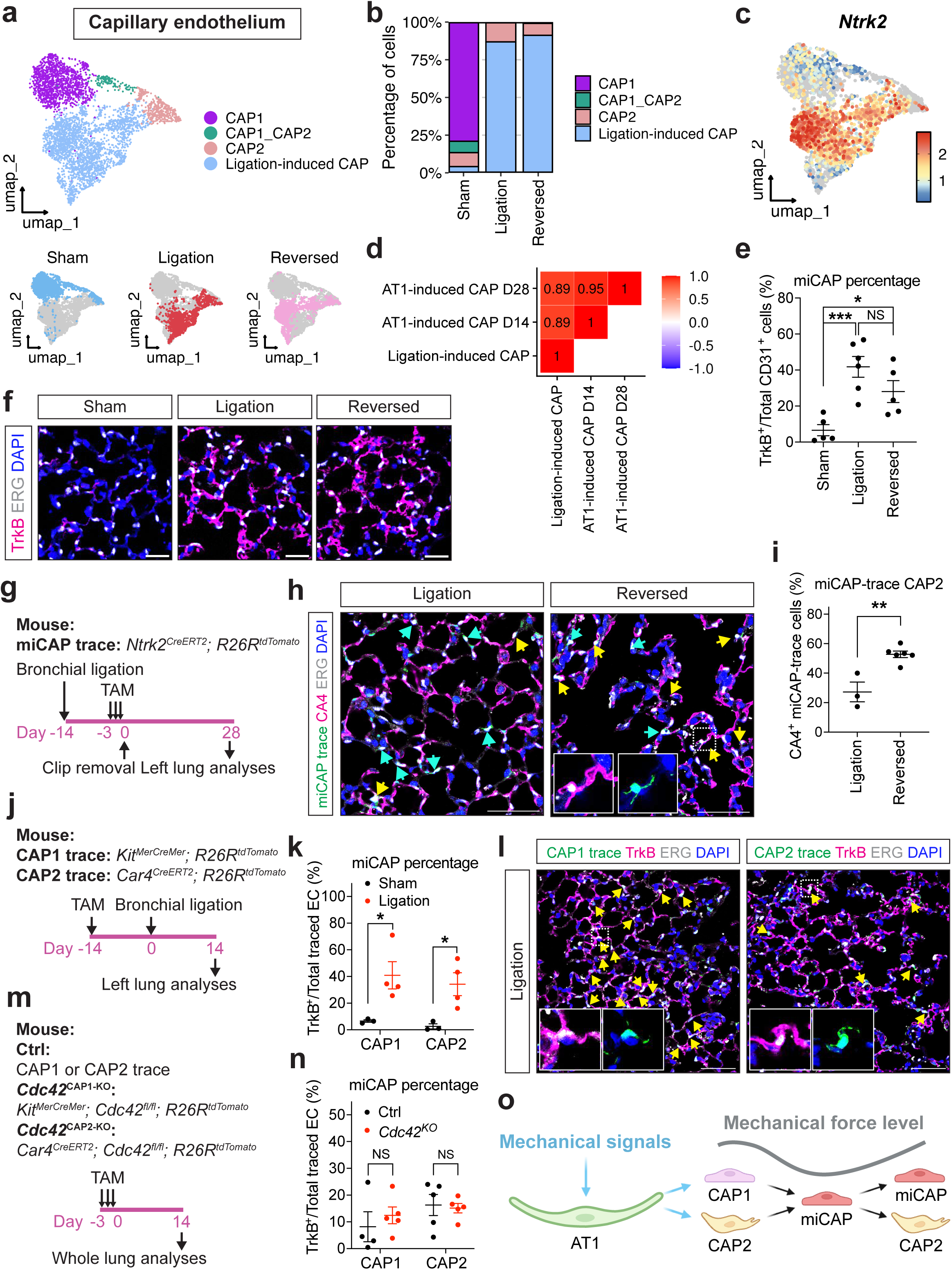
The AT1 mechanosensing-induced CAP state is permanent and functionally distinct. **a.** UMAP plots of CAPs from sham, ligated and reversed mouse lungs. Cells were annotated by cell type or treatment. **b.** A bar plot showing percentage of each cell type in sham, ligated or reversed mouse lungs. **c.** A feature plot showing *Ntrk2* expression in ligation-induced CAPs. **d.** A Spearman correlation showing transcriptional similarity between AT1-induced CAPs and ligation-induced CAPs. **e.** Flow cytometry results showing miCAP percentage in indicated lung tissues (n = 5 or 6, mean ± S.E.M.). Asterisks indicate statistical significance (* *p* value < 0.05, *** *p* value < 0.001, unpaired two-tailed Student’s *t* test). NS: no statistical significance. **f.** Representative IHC images of indicated lung tissues stained with TrkB and ERG, a pan endothelial cell marker (scale bar = 25 μm). **g.** A schematic of miCAP lineage trace experiment. miCAP-trace mice underwent ligation surgery, received three doses of tamoxifen, followed by reversed procedure to restore respiratory motion. Tissues from ligated or reversed miCAP-trace mouse lungs were used for histology. **h.** Representative IHC images of indicated lung tissues stained with the lineage trace marker (tdTomato), a CAP2 marker (CA4), and ERG (scale bar = 50 μm). **i.** Percentage of CA4^+^ miCAP-trace endothelial cells in indicated tissues (n = 3 or 6, mean ± S.E.M.). Asterisks indicate statistical significance (** *p* value < 0.01, unpaired two-tailed Student’s *t* test). **j.** A schematic of CAP lineage trace experiment. CAP1- or CAP2-trace mice were administrated with one dose of tamoxifen, followed by sham or bronchial ligation surgery. Cells and tissues from control or ligated lung were used for flow cytometry or histology, respectively. **k.** Flow cytometry results showing miCAP percentage in indicated lung tissues (n = 3 or 4, mean ± S.E.M.). Asterisks indicate statistical significance (* *p* value < 0.05, unpaired two-tailed Student’s *t* test). **l.** Representative IHC images of CAP1- (left) or CAP2-trace (right) mouse lungs after bronchial ligation stained with the lineage trace marker (tdTomato), TrkB, and ERG (scale bar = 100 μm). **m.** A schematic of genetic attenuation of mechanotransduction in CAPs. Control CAP-trace mice or those with a *Cdc42^fl/fl^* allele were administrated with three doses of tamoxifen. Cells from control or *Cdc42*^CAP-KO^ lung were used for flow cytometry. **n.** Flow cytometry results showing miCAP percentage in indicated lung tissues (n = 4 or 5, mean ± S.E.M.). NS: no statistical significance. **o.** A model depicting miCAP cell origin and fate change in response to mechanical forces.

To investigate the cell of origin for the miCAPs, we utilized two previously validated lineage trace mouse lines, *Kit^MerCreMer^; R26R^tdTomato^* and *Car4^CreERT2^; R26R^tdTomato^* to label CAP1s and CAP2s, respectively, and track their cell state changes upon attenuated respiratory movements (**Fig. 4j**)^17^. CAP1- and CAP2-trace mice were given tamoxifen to initiate lineage tracing, followed by sham or bronchial ligation surgery (**Fig. 4j**). Flow cytometry and IHC analyses indicated that both CAP1 and CAP2 lineage can give rise to miCAPs upon inhibition of respiratory motion (**Fig. 4k-l and Extended Data Fig. 9a**).

To determine whether cell intrinsic mechanotransduction in CAP1 or CAP2 caused their state switch to miCAPs, mouse lines were created to block intrinsic mechanotransduction in either CAP1 or CAP2 by conditional knockout of *Cdc42* (*Cdc42*^CAP1-KO^ or *Cdc42*^CAP2-KO^, **Fig. 4m**). These knockouts successfully attenuated mechanotransduction in CAPs, as indicated by decreased expression of mechanosignaling target genes including *Ccn1*, *Ccn2* and *Ankrd1* (**Extended Data Fig. 9b**)^24^. However, we did not observe the miCAP state arising from either CAP1 or CAP2 lineage upon cell intrinsic inhibition of mechanotransduction (**Fig. 4n and Extended Data. Fig. 9c**). Together, these findings indicate that this permanent miCAP state is functionally distinct and its emergence is induced by defective mechanotransduction in neighboring AT1 cells.

### AT1-endothelial mechanotransduction requires integrin/TGF-β signaling

To define the mechanisms by which AT1 cells sense and transmit mechanical signals to their adjacent CAPs to induce the miCAP state, we analyzed the signaling pathways enriched in CAPs from *Cdc42*^AT1-KO^ mouse lungs using our scRNAseq data (**Fig. 3h and 5a**). The transforming growth factor β (TGF-β) pathway was significantly upregulated in CAPs in response to *Cdc42*^AT1-KO^ (**Fig. 5a**). Of note, TGF-β signaling was also enriched in CAPs from the ligated mouse lungs (**Extended Data Fig. 3c**). To validate altered TGF-β signaling in CAPs, we isolated CD31^+^ ECs from control or *Cdc42*^AT1-KO^ mouse lungs by fluorescence-activated cell sorting (FACS) and examined phosphorylation of Suppressor of Mothers against Decapentaplegic (SMAD) proteins 2/3 (**Fig. 5b and Extended Data Fig. 10a**). Consistent with TGF-β signaling activation, phosphorylated SMAD2/3 in ECs from *Cdc42*^AT1-KO^ mouse lungs was significantly increased relative to that from control (**Fig. 5b-c**)^29,30^. To assess the necessity of TGF-β signaling in the development of miCAP state, we genetically deleted the gene encoding TGF-β receptor II in mouse lung ECs (*Tgfbr2*^EC-KO^) and examined the emergence of miCAPs after bronchial ligation (**Fig. 5d**)^31^. Attenuation of TGF-β signaling transduction inhibited the emergence of miCAPs, shown by both flow cytometry and tissue immunostaining (**Fig. 5e-f and Extended Data Fig. 10b**), indicating that the development of miCAP state requires intact TGF-β signaling activity in CAPs.

**Figure 5:**
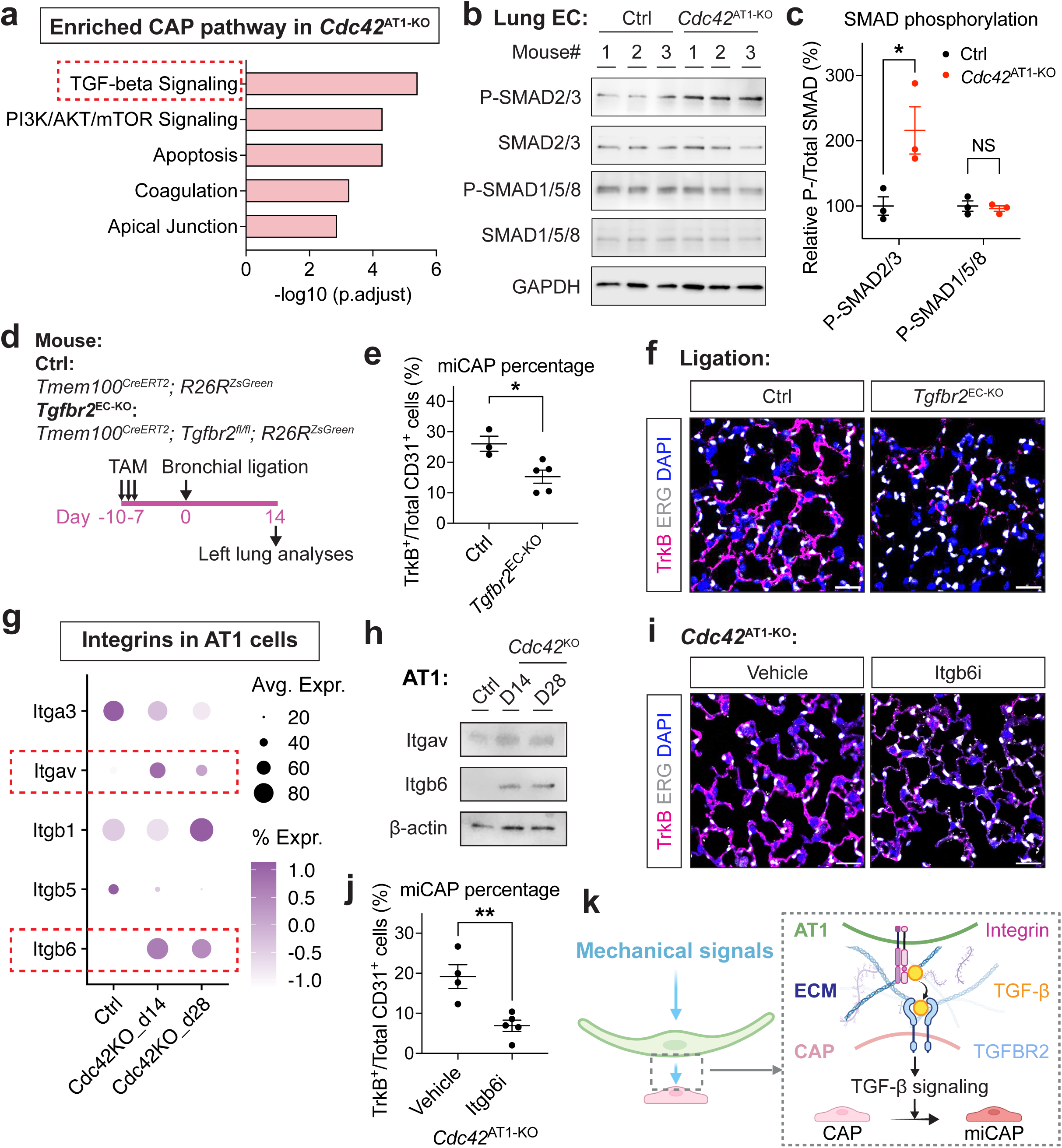
AT1-endothelial mechanotransduction requires integrin/TGF-β signaling. **a.** A pathway enrichment analysis of CAPs from control and *Cdc42*^AT1-KO^ (D14 and D28) mouse lungs. **b.** An immunoblot showing phosphorylation of SMADs in endothelial cells from control and *Cdc42*^AT1-KO^ (D14) mouse lungs. Three independent mice were included in each group. **c.** Quantification of p-SMAD level relative to total SMAD from immunoblots in (**b**) (n = 3, mean ± S.E.M.). An asterisk indicates statistical significance (* *p* value < 0.05, unpaired two-tailed Student’s *t* test). NS: no statistical significance. **d.** A schematic of genetic attenuation of TGF-β signaling in endothelial cells. Control EC-trace mice or those with a *Tgfbr2^fl/fl^* allele were administrated with three doses of tamoxifen, followed by ligation surgery after 7 days. Cells and tissues from ligated control or *Tgfbr2*^EC-KO^ mouse lungs were used for flow cytometry or histology, respectively. **e.** Flow cytometry results showing miCAP percentage in indicated lung tissues (n = 3 or 5, mean ± S.E.M.). An asterisk indicates statistical significance (* *p* value < 0.05, unpaired two-tailed Student’s *t* test). **f.** Representative IHC images of indicated lung tissues stained with TrkB and ERG, a pan endothelial cell marker (scale bar = 25 μm). **g.** A dot plot showing indicated *Integrin* expression in AT1 cells from control and *Cdc42*^AT1-KO^ mouse lungs. Red dashed boxes indicate specific *Integrin* with increased expression after *Cdc42*^AT1-KO^. **h.** An immunoblot showing increased level of Itgav and Itgb6 in AT1 cells from control and *Cdc42*^AT1-KO^ mouse lungs. AT1 cells from three independent mice were pooled and used for protein extraction in each group. **i.** Representative IHC images of lung tissues from *Cdc42*^AT1-KO^ mice in the presence of vehicle or a small-molecule Itgb6 inhibitor administration (scale bar = 25 μm). **j.** Flow cytometry results showing miCAP percentage in indicated lung tissues (n = 4 or 5, mean ± S.E.M.). Asterisks indicate statistical significance (** *p* value < 0.01, unpaired two-tailed Student’s *t* test). **k.** A model depicting integrin/ TGF-β-dependent crosstalk between AT1 cells and CAPs.

TGF-β signaling requires cell surface integrin heterodimers to sequester the latency-associated peptide (LAP) in the latent TGF-β complex, releasing activated TGF-β ligands into the extracellular matrix (ECM) for binding to cognate receptors^29,30,32^. We analyzed the level of integrin genes in AT1 cells using our scRNA-seq data and noted that the expression of both *Itgav* and *Itgb6* were increased in *Cdc42*^KO^ lungs (**Fig. 5g**). Importantly, *Itgb6* is highly expressed in AT1 cells in the mouse and human lung and to a lower level in AT2 cells (**Extended Data Fig. 11a-b**). Immunoblotting using protein from FACS-isolated AT1 cells from control and *Cdc42*^AT1-KO^ mouse lungs confirmed the increased Itgav and Itgb6 protein levels in these cells upon *Cdc42*^KO^ (**Fig. 5h and Extended Data Fig. 10a**). We then determined whether these integrins are responsible for AT1-endothelial mechanotransduction using a small molecule specifically inhibiting Itgb6 heterodimer and its resultant TGF-β activation (Itgb6i)^33^. Compared to vehicle control, the administration of Itgb6i significantly inhibited miCAP emergence in *Cdc42*^AT1-KO^ mouse lungs shown by both flow cytometry and immunostaining (**Fig. 5i-j**). These results together reveal that AT1 cells require Itgb6 integrin-mediated signaling for mechanical signal transmission to their neighboring endothelial cells (**Fig. 5k**).

### AT1 mechanosensing can distinguish human lung disease phenotypes

Many lung diseases are characterized by aberrant lung mechanics, which contributes to abnormal respiratory function^3^. Fibrotic lung diseases, such as idiopathic pulmonary fibrosis (IPF), exhibit increased tissue stiffness that leads to loss of alveolar capacity^3,34^. By contrast, obstructive lung diseases, including chronic obstructive pulmonary disease (COPD), exhibit an emphysematous phenotype resulting in hyperinflated and degenerated alveoli^3,35^. Using scRNAseq datasets from human disease lungs, we generated a mechanosignaling gene expression score using same YAP/TAZ target genes as described above to evaluate the mechanical activity of cells within the alveoli (**Fig. 6a and Extended Data Fig. 12a**)^24,36^. Compared to other cells, AT1 cells showed significant baseline mechanosignaling activity in healthy human lungs, consistent with our mouse data (**Fig. 2b, 6a and Extended Data Fig. 12a**). Notably, AT1 mechanotransduction was increased in COPD lungs but decreased in IPF (**Fig. 6a and Extended Data Fig. 12a**). In addition, we analyzed our recent scRNAseq data from influenza infected mouse lungs and noted a significant decrease in AT1 mechanotransduction in response to influenza lung injury using our mechanoscore (**Extended Data Fig. 12b**)^17^. These findings suggest that AT1 cells are sensitive to both acute viral injury and chronic disease associated changes in lung mechanics.

**Figure 6:**
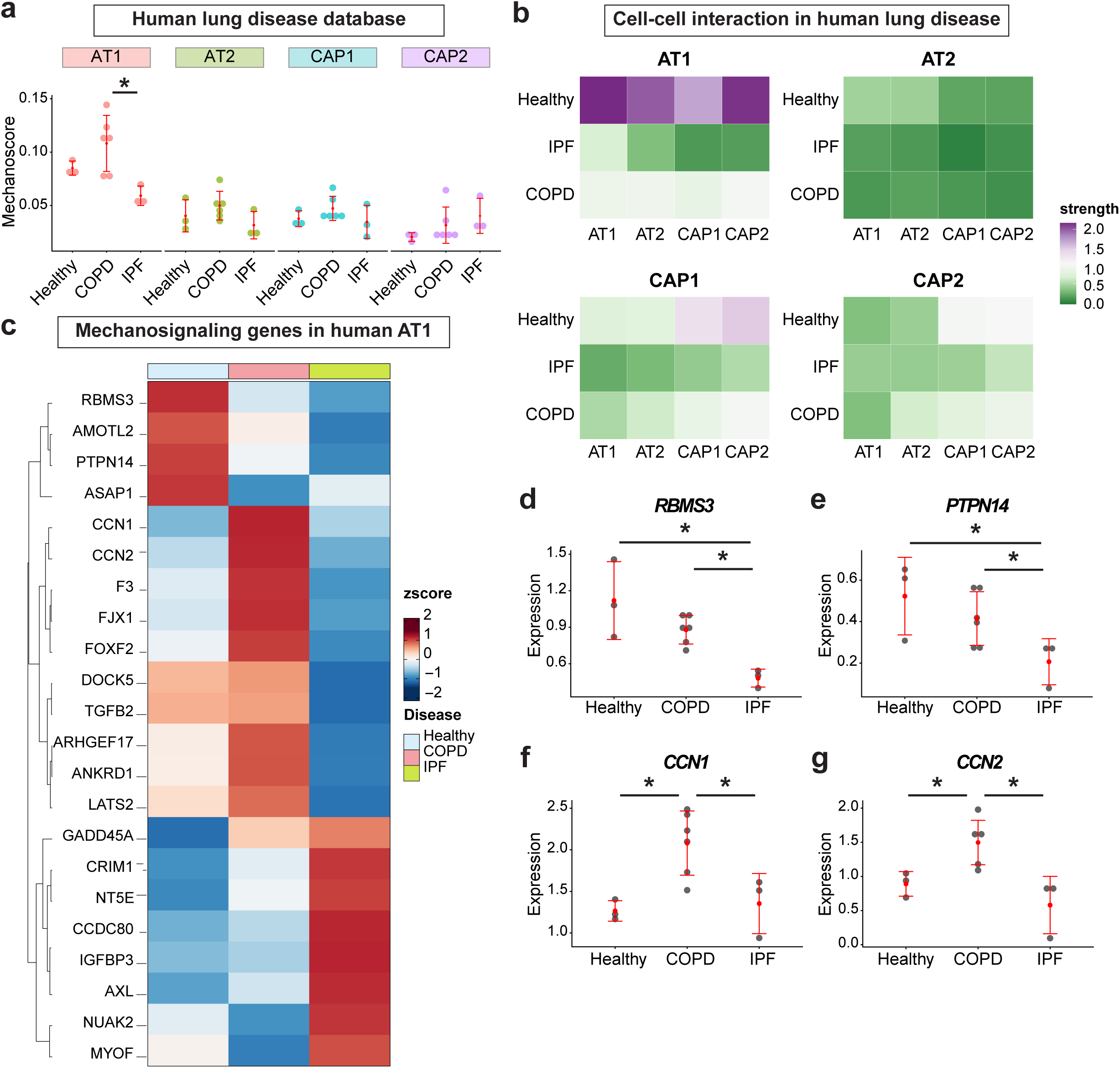
The AT1-driven mechanosensing network is altered in human lung disease. **a.** Mechanoscore of indicated cells from human healthy, COPD, and IPF lung tissues generated using pseudobulk aggregated expression of YAP/TAZ target genes described above in Fig. 2a (median ± S.D.). An asterisk indicates statistical significance (* *p* value < 0.05, Kruskal-Wallis test followed by Dunn test). **b.** CellChat analysis showing communication strength between indicated cell types from human healthy, COPD, and IPF lung tissues. **c.** A heatmap showing expression of YAP/TAZ target genes used for mechanoscore in AT1 cells from human healthy, COPD, and IPF lung tissues. **d-g.** Pseudobulk aggregated expression of indicated genes in AT1 cells from human healthy, COPD, and IPF lungs (median ± S.D.). An asterisk indicates statistical significance (* *p* value < 0.05, Kruskal-Wallis test followed by Dunn test).

We further determined the communication between AT1 cells and their niche neighbors by CellChat analysis using human disease scRNAseq data^36^. AT1 cells showed a significant interaction with their niche neighbors in healthy lungs (**Fig. 6b**). However, in both human COPD and IPF lungs, AT1-driven intercellular communication was greatly reduced, indicating that AT1 cells may require an appropriate level of mechanical load to maintain crosstalk with adjacent cells (**Fig. 6b**). Interestingly, expression of mechanosignaling target genes showed distinct disease-associated patterns (**Fig. 6c**). Expression of a subset of genes, such as *RBMS3* and *PTPN14,* were dramatically downregulated in AT1 cells from IPF samples (**Fig. 6c-e**). Other mechanosignaling genes including *CCN1* and *CCN2* were remarkably increased in AT1 cells from COPD lungs, whereas these genes showed minimal changes in those from IPF (**Fig. 6c, f-g**). These results suggest that AT1 cells rely on different transcriptional programs to mediate their mechanosensing in fibrotic versus emphysematous diseases. Taken together, our findings highlight that AT1 cells are a hub of alveolar mechanotransduction, propagating mechanical cues during respiratory movements to maintain overall homeostasis of the lung alveolar niche.

## Discussion

Biophysical forces impact cell and tissue behaviors in organs such as the lungs. However, which cells are sensitive across the various regions of the lung and how such forces are interpreted and propagated is not understood. Leveraging a reversible bronchial ligation model, we have now demonstrated that respiration-mediated mechanical forces are essential for maintaining multi-cellular homeostasis in the lung alveolus. These studies show that the AT1 lineage is a hub of mechanical signaling in the alveolus, with AT1-specific blockade of mechanotransduction transcriptionally altering neighboring cell lineages including CAPs. The AT1 mechanosensing-induced changes in CAPs are significant and persistent, and are driven in part through integrin/TGF-β signaling resulting from aberrant AT1 mechanosignaling. Mechanotransduction also occurs in human AT1 cells and is altered in disease, and these changes can phenotypically distinguish fibrotic versus emphysematous diseases. Thus, our study defines the niche-dependent role for mechanotransduction in the lung alveolus and reveals that changes in specific cell behavior and response are altered in chronic human lung diseases.

Aberrant lung mechanics is a hallmark of various human lung diseases^3^. Chronic lung diseases such as COPD and IPF are characterized by alterations in tissue elasticity and stiffness, respectively^34,35^. In addition, in human patients, forced ventilation of the lungs in acute diseases such as acute respiratory distress syndrome (ARDS) can lead to severe lung damage due to changes in mechanical load^37^. The bronchial ligation model may lead to a better understanding of mechanisms driven by deregulated lung mechaosignaling and provide insights on how aberrant mechanics impact lung tissue structure and function in lung diseases. While genetic models of mechanotransduction have led to important discoveries in lung homeostasis and function, most of these models represent irreversible modulation in mechanotransduction and thus cannot be utilized to assess the alterations that occur upon resolution of such inhibitory impulses^4,8,25^. The development of the reversible bronchial ligation model has allowed us to study cellular changes in response to release of lung motion blockade. The changes in cell transcriptome and state we have observed suggest that even if lung motion and tissue architecture can be normalized, persistent changes in cell state and behavior can occur, which may further impact tissue responses to therapeutic interventions and subsequent insults and injuries. Moreover, clinical treatments of various lung diseases involve interventions in lung mechanical load, such as mechanical ventilation^38^. The reversible bronchial ligation model may also help understand whether and how such temporarily altered lung mechanics may impact tissue structure and function in the long term.

Altered mechanosignaling in AT1 cells impacts the fate of multiple neighboring cell types including endothelial cells. Blocking AT1 cell mechanotransduction induces a *Ntrk2^+^/CD74^+^* CAP state (miCAP) that is also observed in our bronchial ligation model. Previous work has also revealed a transcriptionally similar endothelial cell state that emerges after acute viral lung injuries and chronic lung fibrosis^17,19,39,40^. suggesting the existence of miCAPs may arise from dysfunctional AT1 mechanosensing during both acute and chronic diseases states. Compared to *Ntrk2*-negative CAPs, miCAPs spontaneously respond to normalization of respiration-mediated mechanical forces in our reversible bronchial ligation model by re-entering the cell cycle and differentiating into CAP2s, indicating these cells can act as a potential progenitor during endothelial regeneration after mechanical insults.

How mechanical forces are propagated between cells within a tissue niche has remained poorly understood due to the complexities of studying multi-cellular biophysical force changes *in vivo*. The induction of the miCAP state upon loss of mechanical signaling in AT1 cells is dependent on endothelial TGF-β signaling induced by the AT1-specific integrin Itgb6. Multiple studies have implicated integrin signaling including that by Itgb6 heterodimers in fibrotic diseases such as IPF. This has led to the development of therapeutic strategies to treat lung fibrosis through inhibition of integrins that blocks the resultant activation of TGF-β ligands^32,41–45^. This approach is validated by our findings that Itgb6 blockade using a small molecule dramatically inhibited CAP phenotypic changes driven by TGF-β signaling activation. Since *Itgb6* expression is restricted to AT1 cells in both the mouse and human lung, blocking its function could have deleterious effects on certain AT1-dependent lung function. Thus, while pharmacological Itgb6 inhibition can efficiently reduce TGF-β-dependent tissue stiffening, it may also disrupt AT1 mechanosensing-mediated tissue homeostasis and regeneration, leading to unexpected side effects. Therefore, a detailed characterization of integrin function in distinct cell types may need to be revisited as these therapies are developed for IPF and other diseases.

Lung diseases span a range of phenotypic changes in tissue architecture from pulmonary fibrosis which is characterized by increased extracellular matrix (ECM) deposition and stiffing of lung tissue, to emphysematous diseases such as COPD which are marked by loss of alveolar structure and impaired tissue elasticity^3,34,35^. Our analysis reveals aberrant mechanotransduction in human AT1 cells in both IPF and COPD lungs, suggesting AT1 mechanotransduction is sensitive to abnormal tissue compliance. Whether and how AT1 mechanosensing-mediated cell responses play a role in the onset and progression of these diseases requires further investigation. Mechanotransduction in AT1 cells maintains the integrity of ECM through TGF-β signaling and may act as a potential mechanism underlying compromised ECM deposition and function during progressive pulmonary fibrosis and emphysema^28^. Furthermore, AT1 mechanosignaling-driven gene transcription is distinct between IPF and COPD, suggesting that specific transcription program may direct tissue towards fibrotic or emphysematous responses.

Taken together, our studies highlight that AT1 cells propagate biophysical signals induced by respiratory movements to maintain overall lung homeostasis. Dysregulated AT1 mechanosensing leads to significant changes in fate and behavior of alveolar cells that are essential for lung structure and function. In addition, AT1 mechanosensing is altered in both acute and chronic lung diseases and can help phenotypically differentiate between fibrotic versus emphysematous diseases. Future studies characterizing mechanosensing-induced tissue responses in complex organs such as the lung will be critical to understand the mechanisms underlying onset and progression of various diseases and could provide directions for the development of new therapeutics.

## Methods

### Human subjects

All human samples used in this study were obtained following an established protocol (Prospective Registry of Outcomes in Patients Electing Lung Transplantation (PROPEL)) approved by University of Pennsylvania Institutional Review Board, as previously reported^36^. Informed consent was performed in accordance with institutional and NIH procedures. The institutional review board of the University of Pennsylvania approved this study, and all patient information was deidentified before use.

### Mouse studies

All mouse experiments were performed following the protocols of the University of Pennsylvania Institutional Animal Care and Use Committee. Adult mice (8-12 weeks old) were used in all the experiments. Cre recombination was initiated by a single dose of tamoxifen (200 mg/kg Sigma-Aldrich) for lineage tracing and 1 dose of tamoxifen (200 mg/kg, Sigma-Aldrich) daily for 3 consecutive days for conditional gene knockout, unless otherwise indicated. The small-molecule Itgb6 inhibitor EMD527040 (Tocris Bioscience Cat#7508) was dissolved in 10% DMSO in saline and administrated intraperitoneally at 20 mg/kg every other day.

Mouse lines used in this study were previously validated and the strains are listed below:

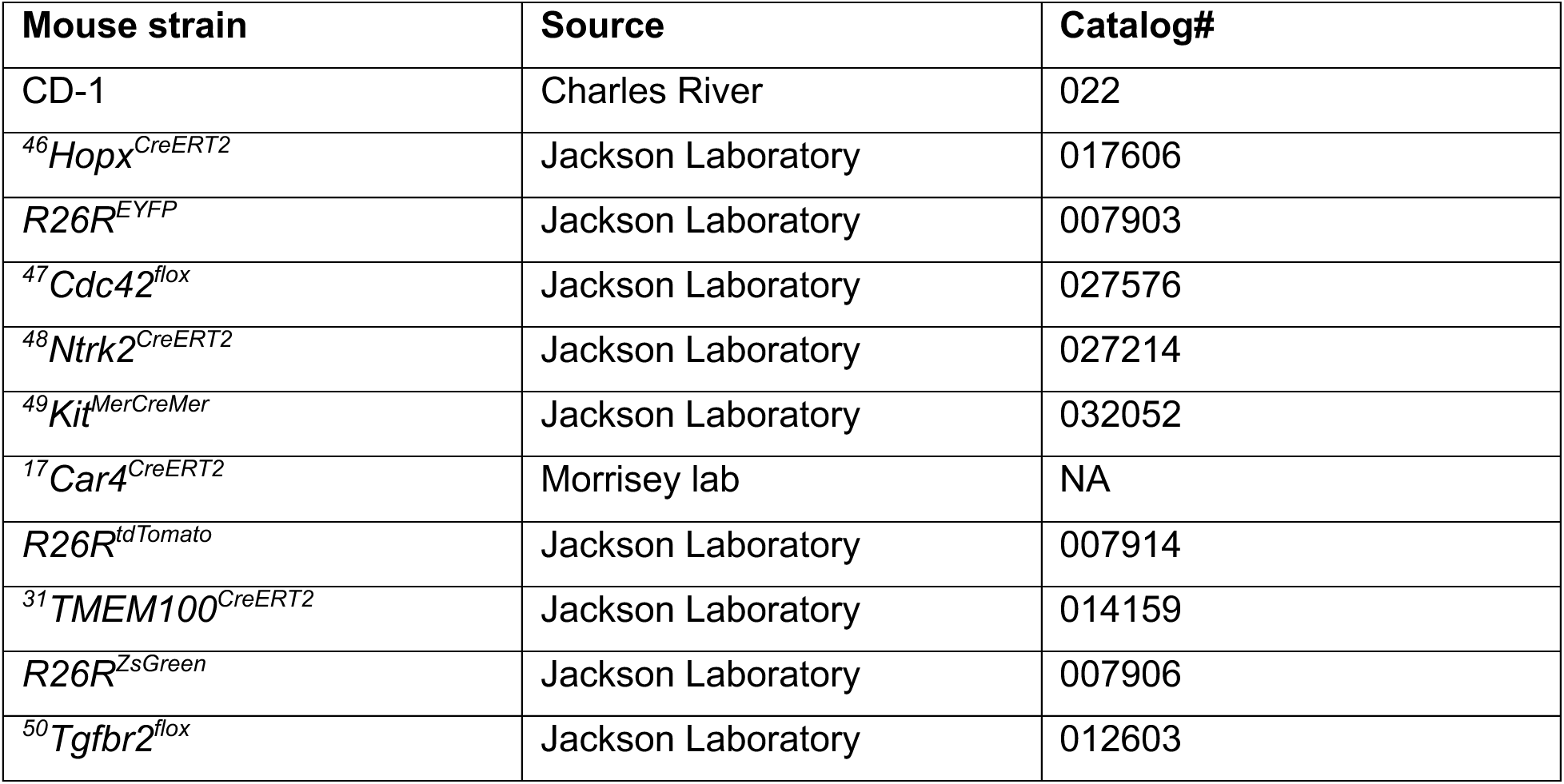

### Reversible bronchial ligation

Mice were anesthetized with isoflurane and intubated for ventilation using a MiniVent ventilator (Harvard apparatus; tidal volume of 10 μL per gram mouse body weight, respiratory rate of 150 breaths per minute). The thoracic cavity was exposed by incising 1 cm on the left lateral side of the skin, followed by a 0.5 cm incision at the third left intercostal space. The left main bronchus was clipped using a customized micro vascular clamp (Fine Science Tools Cat#00396-01, jaw dimension = 2 x 1 mm) with a micro clamp applying forceps (Fine Science Tools Cat#00071-14). For reversal, mice were anesthetized and intubated following the same protocol. An incision was made at the fourth left intercostal space to expose thoracic cavity, followed by removal of the clip.

### Micro-computed tomography (uCT)

uCT images were generated using a X-Cube CT scanner (Molecubes) located at the Small Animal Imaging Facility at Penn Medicine. Live mice were anesthetized with isoflurane and images were acquired using a respiratory gating approach. The settings for image acquisition were as follows: reconstructed voxel size: 100 um, X-Ray tube voltage: 50 kVp, X-Ray tube current: 440 uA, exposure time: 32 ms, type of filter used: 0.8 mm aluminum filter, degree of rotation stepping: 0.75 degrees (480 total steps in a 360 degree scan).

### Histology

For tissue preparation, mice were euthanized by CO_2_ inhalation and the lungs were perfused with PBS, followed by inflation with 2% Paraformaldehyde (Thermo Fisher Scientific) at a constant pressure of 25 cm H_2_O. Tissues were fixed at 4°C overnight and dehydrated in a series of ethanol concentration gradients. After dehydration, tissues were embedded in paraffin and sliced into 6 μm thick sections.

Hematoxylin and eosin (H&E) staining was performed as previously described^17,36^. For immunostaining, the following antibodies were used on paraffin tissue sections:

| Antibody | Source | Catalog# |
| --- | --- | --- |
| Goat polyclonal anti-TrkB | Novus Biologicals | AF1494 |
| Mouse monoclonal anti-ERG | Abcam | ab214341 |
| Rabbit monoclonal anti-ERG | Abcam | ab92513 |
| Rabbit polyclonal anti-RFP | Rockland | 600-401-379 |
| Rat monoclonal anti-LAMP3 | Novus Biologicals | DDX0191P-100 |
| Mouse monoclonal anti-HOPX | Santa Cruz | sc-398703 |
| Rabbit monoclonal anti-NKX2.1 (TTF1) | Abcam | ab76013 |
| Goat polyclonal anti-CA4 | Novus Biologicals | AF2414 |

### Imaging and image analysis

H&E stained lung sections were tile-scanned at 4x using a Nikon Eclipse Ni series microscope. Fluorescent images were acquired at 20x and 40x using z-stacks on a Zeiss LSM 710 or Leica Stellaris 5 laser scanning confocal microscope. Cell counting analysis was performed using FIJI.

### Single cell isolation, flow cytometry, and FACS

Lungs were harvested and digested into single cell suspensions as previously described^17,36^. For single cell dissociation from ligated mouse lungs, lungs were perfused, inflated with PBS, followed by inflation with digestion buffer supplemented with collagenase I, dispase, and DNase I prior to the removal of lobes.

For flow cytometry, cells were stained with combination of antibodies for 30 min on ice before subjected to a flow cytometer. For FACS, lung cells were incubated with mouse CD45 MicroBeads magnetic beads (Miltenyi Biotec Cat#130-052-301) and the CD45^+^ population was depleted as per the manufacturer’s protocol. CD45^-^ cells were then incubated with CD31 or/and CD326 antibodies and subjected to FACS for isolation of specific cell populations.

Antibodies used and dilutions are listed below:

| **Antibody (Dilution)** | **Source** | **Catalog#** |
| --- | --- | --- |
| CD45-PerCP-Cy5.5 (1:200) | Thermo Fisher | 45-0451-82 |
| CD45-PE-Cy7 (1:200) | Thermo Fisher | 25-0451-82 |
| CD45-APC-Cy7 (1:200) | BioLegend | 103116 |
| CD31-PE-Cy7 (1:200) | Thermo Fisher | 25-0311-82 |
| CD31-PE (1:200) | BioLegend | 102508 |
| CD326-PE-Cy7 (1:200) | Thermo Fisher | 25-5791-80 |
| TrkB-Biotin (1:100) | R&D systems | BAF1494 |
| Streptavidin-APC (1:200) | Thermo Fisher | 17-4317-82 |

### EdU proliferation assay

5-Ethynyl-2’-deoxyuridine (Santa Cruz Biotechnology Cat#sc-284628B) was dissolved in tap water at a concentration of 2 g/L, filtered through a 0.2 μm filter, and stored at 4°C. To create a drinking water solution, this stock solution was diluted to a final concentration of 0.2 g/L in tap water and sterile filtered. Mice consumed EdU water over the time course indicated. EdU incorporation was detected using the Click-iT™ Plus EdU Alexa Fluor™ 488 Flow Cytometry Assay Kit (Thermo Fisher #C10632) according to the manufacturer’s protocol.

### Immunoblotting alaysis

Total protein was extracted from FACS sorted lung endothelial cells or AT1 cells from indicated animals using SDS sample buffer (BioRad) supplemented with 2-Mercaptoethanol (Sigma) and a cocktail of protease and phosphatase inhibitors (Thermo). Samples were boiled for 5 min, followed by SDS polyacrylamide gel electrophoresis. Proteins were then transferred to a nitrocellulose membrane and immunoblotted. Primary and secondary antibodies used are listed as below:

| **Antibody (Dilution)** | **Source** | **Catalog#** |
| --- | --- | --- |
| Rabbit monoclonal P-SMAD2/3 (1:1000) | Cell Signaling Technology | 8828 |
| Rabbit monoclonal SMAD2/3 (1:1000) | Cell Signaling Technology | 8685 |
| Rabbit polyclonal P-SMAD1/5/8 (1:1000) | Millipore Sigma | AB3848-I |
| Rabbit polyclonal SMAD1/5/8/9 (1:1000) | Abcam | ab13723 |
| Rabbit monoclonal GAPDH (1:1000) | Cell Signaling Technology | 8884 |
| Rabbit monoclonal Itgav (1:1000) | Abcam | ab179475 |
| Rabbit monoclonal Itgb6 (1:1000) | Abcam | ab187155 |
| Rabbit monoclonal β-actin (1:1000) | Cell Signaling Technology | 4970 |
| Donkey anti-mouse HRP (1:10000) | Jackson ImmunoResearch | 715-035-150 |
| Donkey anti-rabbit HRP (1:10000) | Jackson ImmunoResearch | 711-035-152 |

### Quantitative RT-PCR

Total RNA was extracted from sorted CAP-trace cells using a RNeasy Mini Kit (Qiagen Cat#74104), reverse transcribed to cDNA using a SuperScriptTM IV kit (Invitrogen Cat#18091050), and used for quantitative PCR reaction on a QuantStudio 7 PCR System (Applied Biosystems) using specific Taqman Probes (Thermo Fisher) or SYBR-Green primers (Integrated DNA Technologies) as listed below:

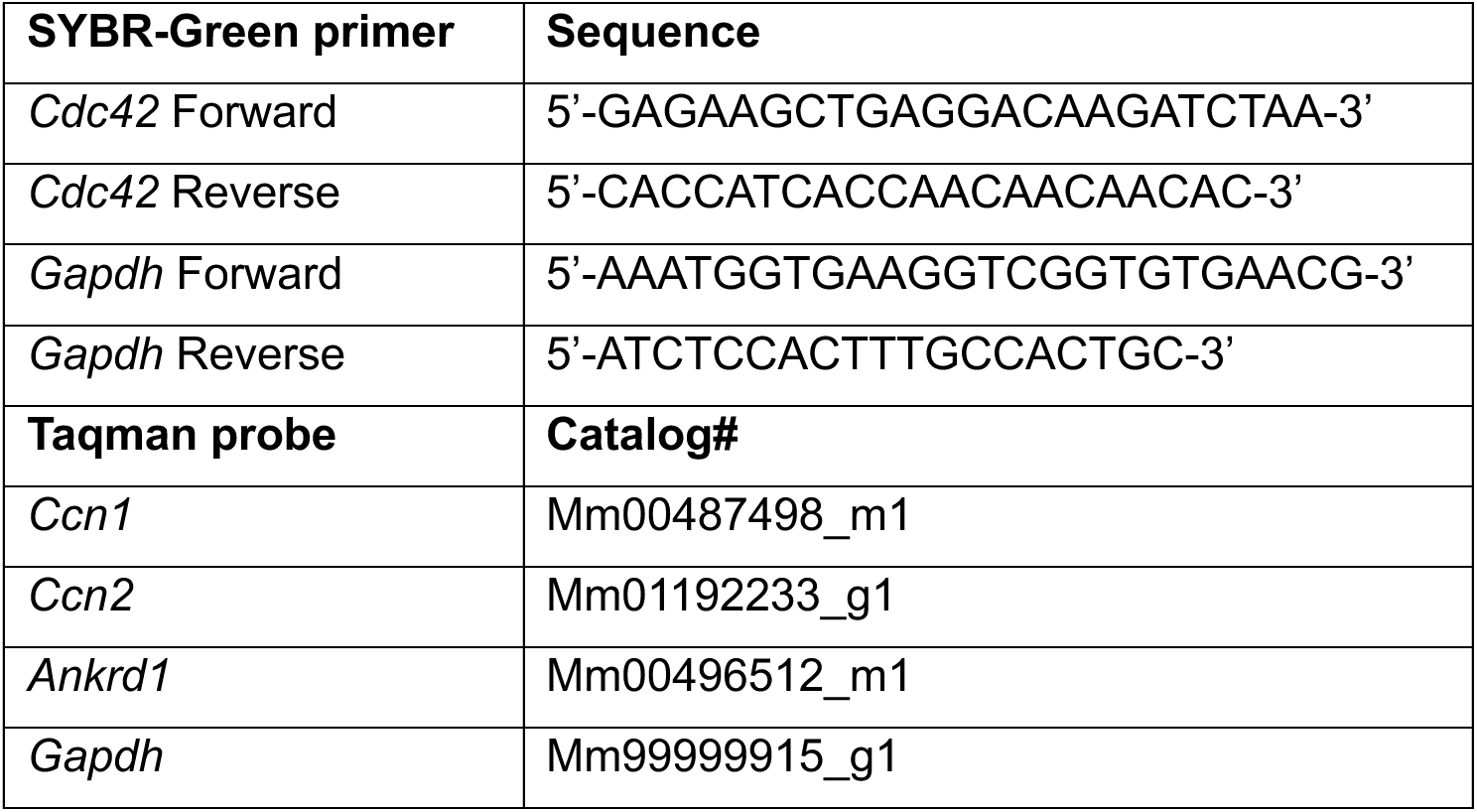

### scRNAseq and analysis

To create samples for scRNAseq that adequately represented both immune and non-immune compartments, we used magnetic activated cell sorting (MACS) to isolate CD45^+^ and CD45^-^ cells and recombined them at a ratio of 85% CD45- cells to 15% CD45+ cells. Cells were counted using the Countess II FL Automated Cell Counter (Thermo Fisher) and loaded at a concentration of 800-1000 cells/mL onto a 10X Chromium Next Gem Controller (10X Genomics) following the manufacturer’s instructions. cDNA and sequencing libraries were prepared using Chromium Next GEM Single Cell 3’ kit v3.1. cDNA and library quality control were performed using an Agilent 2100 Bioanalyzer using the Bioanalyzer High Sensitivity DNA kit (Agilent #5067-4626). Libraries were then sequenced on an Illumina Novaseq 6000 instrument.

Read Alignment, UMI Processing, and Gene Quantification: Raw sequencing data (BCL files) were converted to FASTQ format using bcl2fastq. Reads were aligned to the GRCm39 mouse reference genome using STARsolo (STAR v2.7.11b)^51^ for alignment, barcode and UMI correction, and gene-level quantification in single-cell mode. STARsolo was run with 10x Genomics Cellranger chemistry presets and configured to output both exonic gene counts (--soloFeatures GeneFull). Only barcodes classified as valid cells by STARsolo’s Emptydrops implementation.

Ambient RNA Correction: To reduce contamination from ambient RNA generated by lysed cells during tissue dissociation, we applied SoupX (v1.6.0)^52^ to each sample independently. For each library, SoupX estimated the proportion of background RNA by identifying expression signatures enriched in empty droplets relative to putative cell-containing droplets. The corrected count matrix was obtained using the adjustCounts() function with sample-specific contamination fractions estimated via the autoEstCont() workflow. Post-correction matrices were manually inspected to ensure appropriate removal of canonical ambient signals (e.g., mitochondrial transcripts, hemoglobin genes and ribosomal RNAs). Only contamination-adjusted matrices were used for downstream analysis.

Quality Control and Filtering: Cells with fewer than 200 detected genes, cells exceeding three median absolute deviations (MADs) above the gene-count distribution, cells with a log10GenesPerUMI less than 80% and cells with more than 20% mitochondrial RNA content were excluded. Potential doublets were identified using Scrublet and excluded. All remaining cells were retained for downstream analysis. For experiments that included mixed sexes, X- and Y-linked genes were removed from the variable feature set prior to PCA to prevent sex-specific effects from dominating downstream analyses. Non-linear dimensionality reduction was performed using Uniform Manifold Approximation and Projection (UMAP), and Louvain graph-based clustering was applied, both using the top 30 principal components.

Marker genes for each cell type were identified using the FindAllMarkers command using the RNA assay within Seurat. Correlation analysis was performed in R and correlation plots made using ggcorrplot. Gene set enrichment analysis (GSEA) of gene ontology (GO) was performed using the ‘clusterProfiler’ package (DOI: 10.18129/B9.bioc.clusterProfiler). Mechanoscore gene signatures were obtained from a previously published YAP/TAZ target gene set^24^. Gene signature activity scores were computed using the UCell R package, which implements a rank-based method to quantify gene set enrichment at the single-cell level^53^.

Cell–cell communication analysis was performed using CellChat v2, which infers intercellular signaling networks by integrating known ligand–receptor interactions with single-cell transcriptomic data^54^. For this study, CellChat analysis was restricted to a biologically relevant subset of cell types derived from the human disease samples. All analyses were carried out following the standard CellChat workflow, including data preprocessing, identification of overrepresented ligand–receptor pairs, inference of communication probabilities, and visualization of network-level signaling patterns.

To minimize technical and biological variation, we used a pseudobulk framework in which single-cell counts were aggregated at the donor–cell type level. For each annotated cell type, all cells from the same donor were summed to generate a donor-level expression profile. Pseudobulk matrices were constructed using the Seurat2pb function implemented within the edgeR package^55^. Heatmaps were constructed using the Complexheatmap package in R.

### Statistical analysis

In all mouse experiments, statistical significance was determined by an unpaired two-tailed Student’s *t* test between two groups using Graphpad Prism 10. Statistical analysis of scRNA seq data was performed in R. Details of statistical analysis used for each experiment are indicated in the figure legends.

## Acknowledgements

We thank the Flow Cytometry Core Laboratory at the Children’s Hospital of Philadelphia, the Cell and Developmental Biology Microscopy Core at the University of Pennsylvania (UPenn), and the Rodent Cardiovascular Phenotyping Core at the Cardiovascular Institute of UPenn, for their technical assistance. This research was supported by the National Institutes of Health (R01HL164929, R01HL152194, R01HL132999, R01HL162683, R01HL168803 to E.E.M.).

## Author contributions

Conceptualization: C.S. and E.E.M. Methodology: C.S., M.P.M., S.E.S., G.Z., J.D.P., U.V.C., D.L.J., M.K., Y.Y., S.Z., S.L., H.H., and A.L. Investigation: C.S., M.P.M., S.E.S., G.Z., and J.D.P. Visualization: C.S., M.P.M., J.D.P., and E.E.M. Supervision: M.C.B. and E.E.M. Writing: C.S. and E.E.M.

## Data availability

All newly generated mouse scRNAseq data have been deposited into GEO (GSE310385).

## Extended data figure legends

**Extended Data Figure 1:**
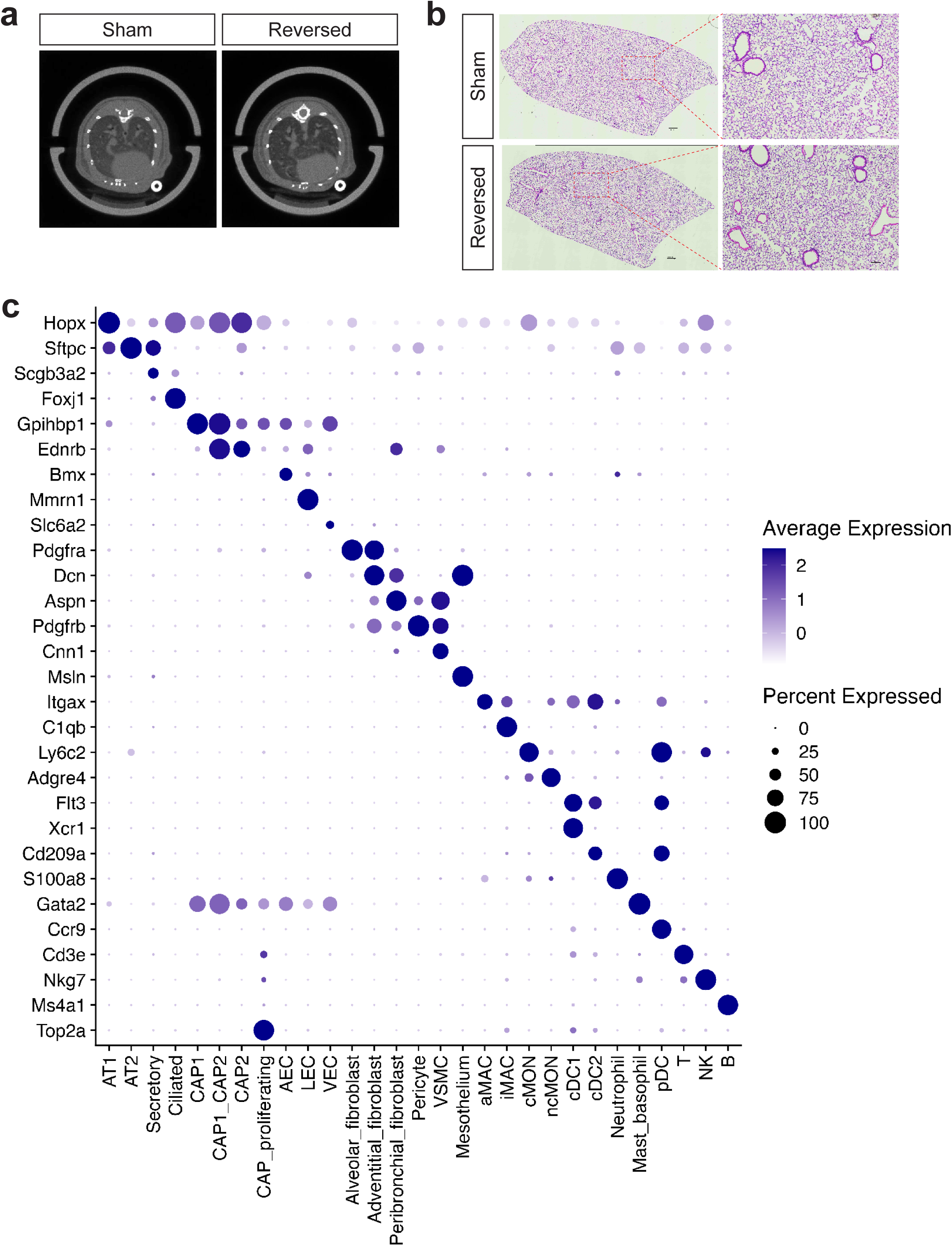
Reversed surgery rescues motion blockade from bronchial ligation. **a.** Representative micro-CT scan images showing restored lung inflation after clip removal in mouse ligated lungs. A schematic of experiment is shown in Fig. 1b. **b.** Representative H&E images of indicated mouse lungs. A schematic of experiment is shown in Fig. 1b. **c.** A dot plot showing marker gene expression in each cell type from the reversible bronchial ligation scRNAseq dataset. A schematic of experiment is shown in Fig. 1b and a UMAP is shown in Fig. 1c.

**Extended Data Figure 2:**
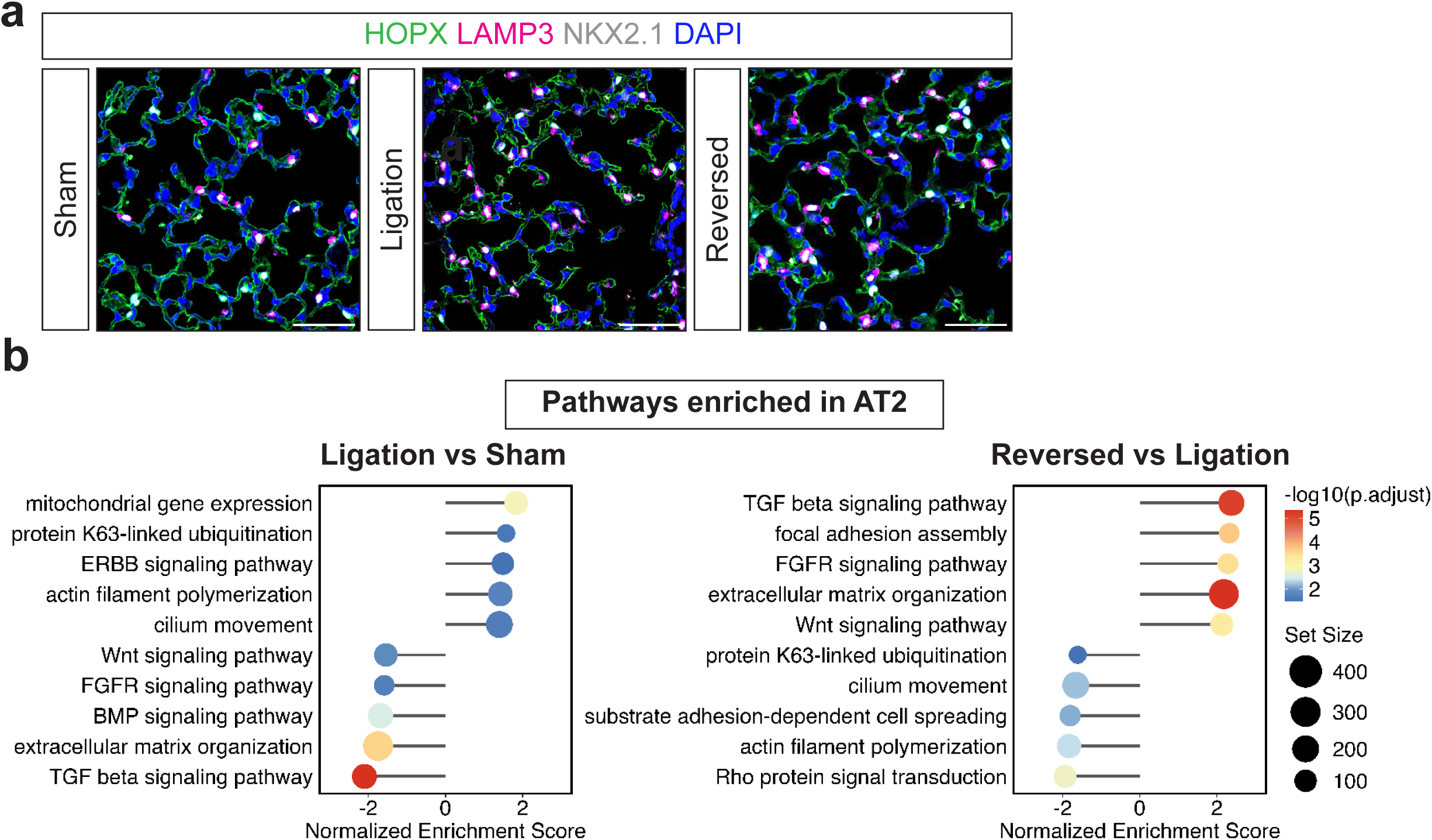
Respiration-mediated mechanical forces control alveolar epithelial composition. **a.** Representative IHC images of indicated lung tissues stained with HOPX (AT1 cell marker), LAMP3 (AT2 cell marker) and NKX2.1 (pan epithelial cell marker) (scale bar = 50 μm). A quantification of AT1/AT2 ratio in each condition is shown in Fig. 1g. **b.** Gene set enrichment analysis results showing altered pathways in AT2 cells from indicated mouse lungs.

**Extended Data Figure 3:**
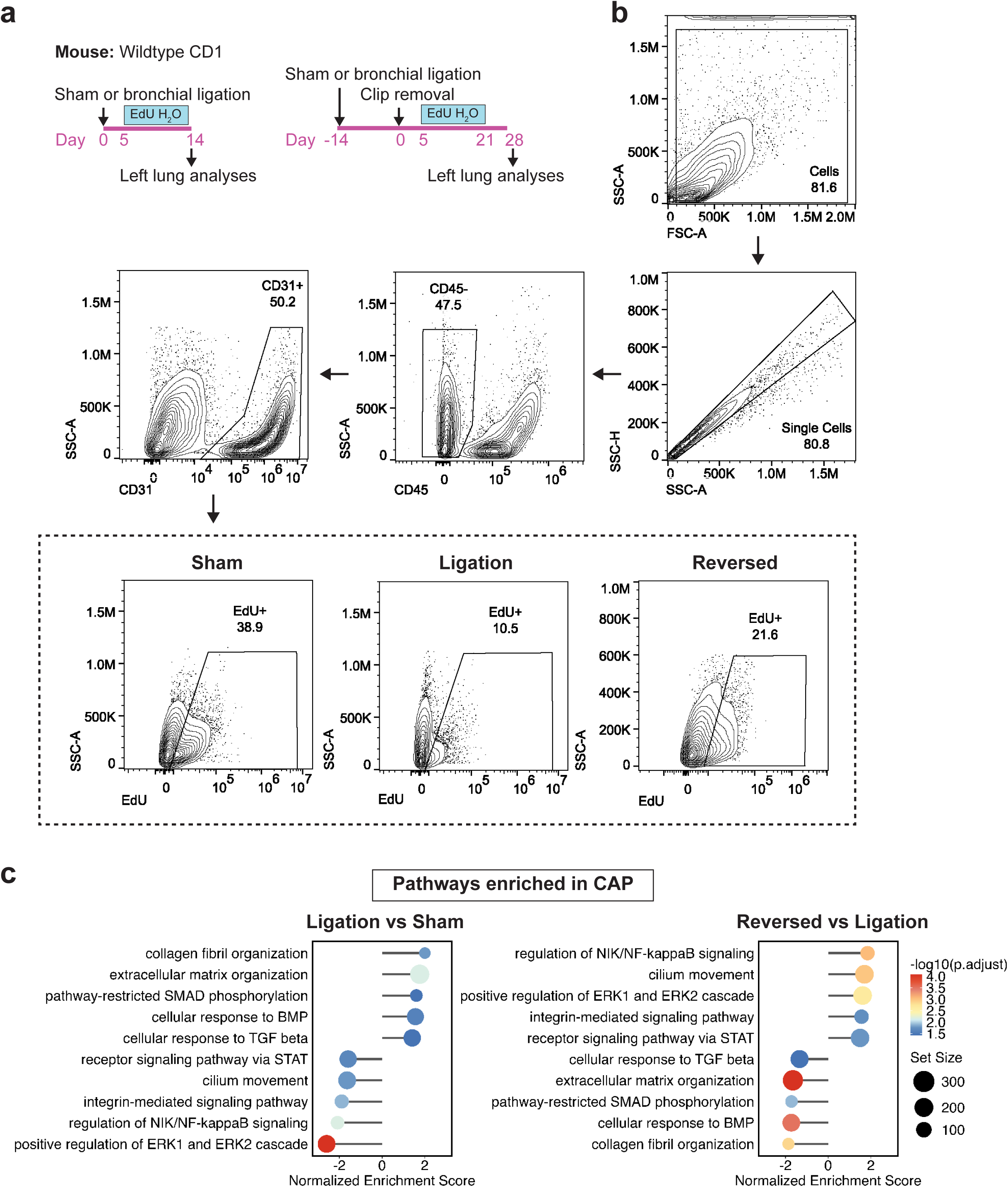
Respiration-mediated mechanical forces control alveolar endothelial cell proliferation. **a.** A schematic of flow cytometry experiment determining proliferation in lung endothelial cells (ECs). EdU water (0.2 g/L) was provided to mice during indicated time periods. Cells from wildtype mouse after sham, ligation or reversed surgery were isolated and used for flow cytometry. **b.** Representative gating strategies of flow cytometry detecting EdU^+^ ECs. A quantification of EdU^+^ ECs in each condition is shown in Fig. 1h. **c.** Gene set enrichment analysis results showing altered pathways in capillary endothelial cells (CAPs) from indicated mouse lungs.

**Extended Data Figure 4:**
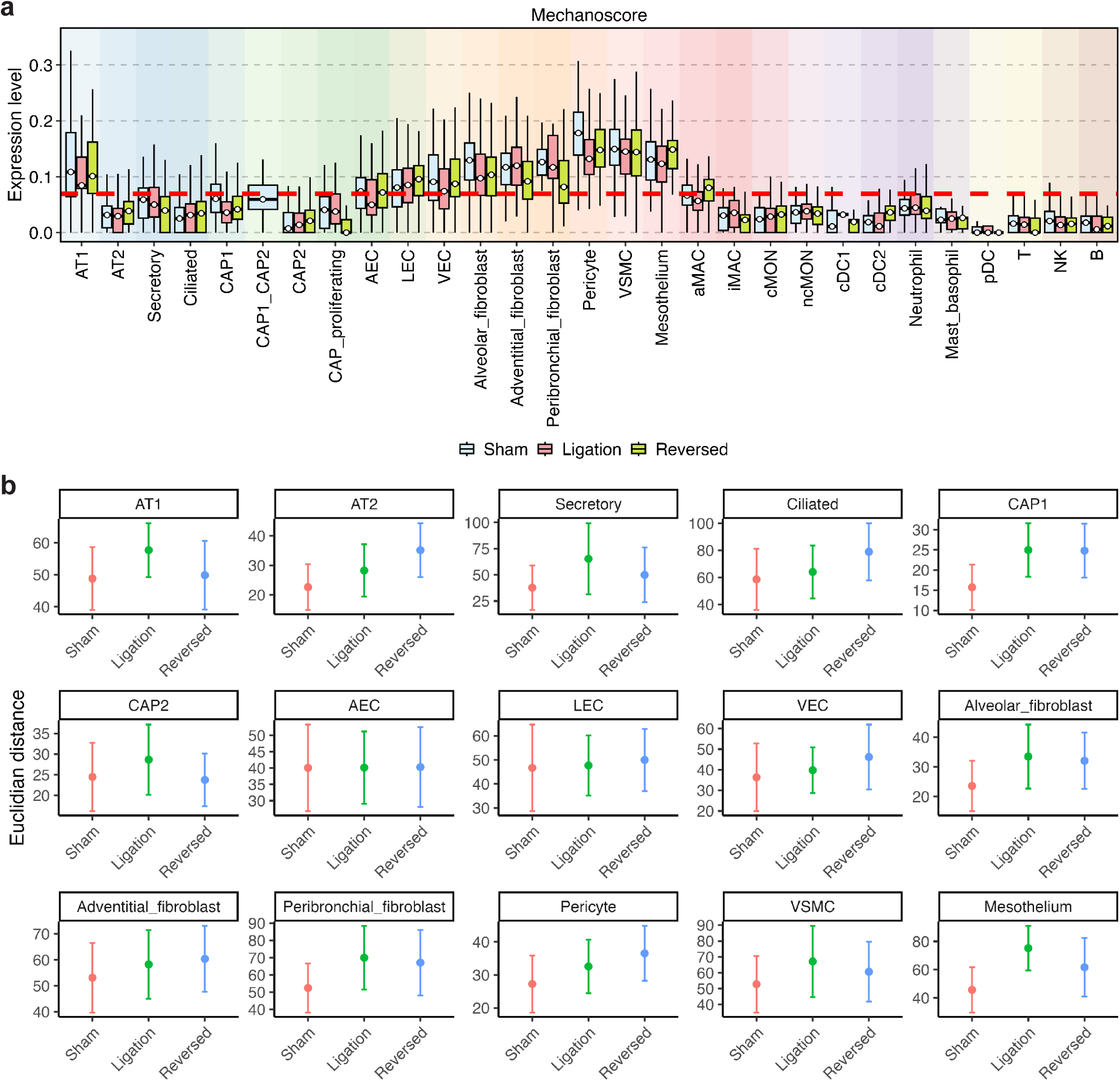
Respiration-mediated mechanical forces control gene transcription in lung cells. **a.** Mechanoscore of each cell type in the distal lung after sham, ligation or reversed surgery. Red dashed line indicates the mean of mechanoscore across all cells at baseline. **b.** Euclidian distance indicating cellular transcriptional similarity across treatments in indicated cell types.

**Extended Data Figure 5:**
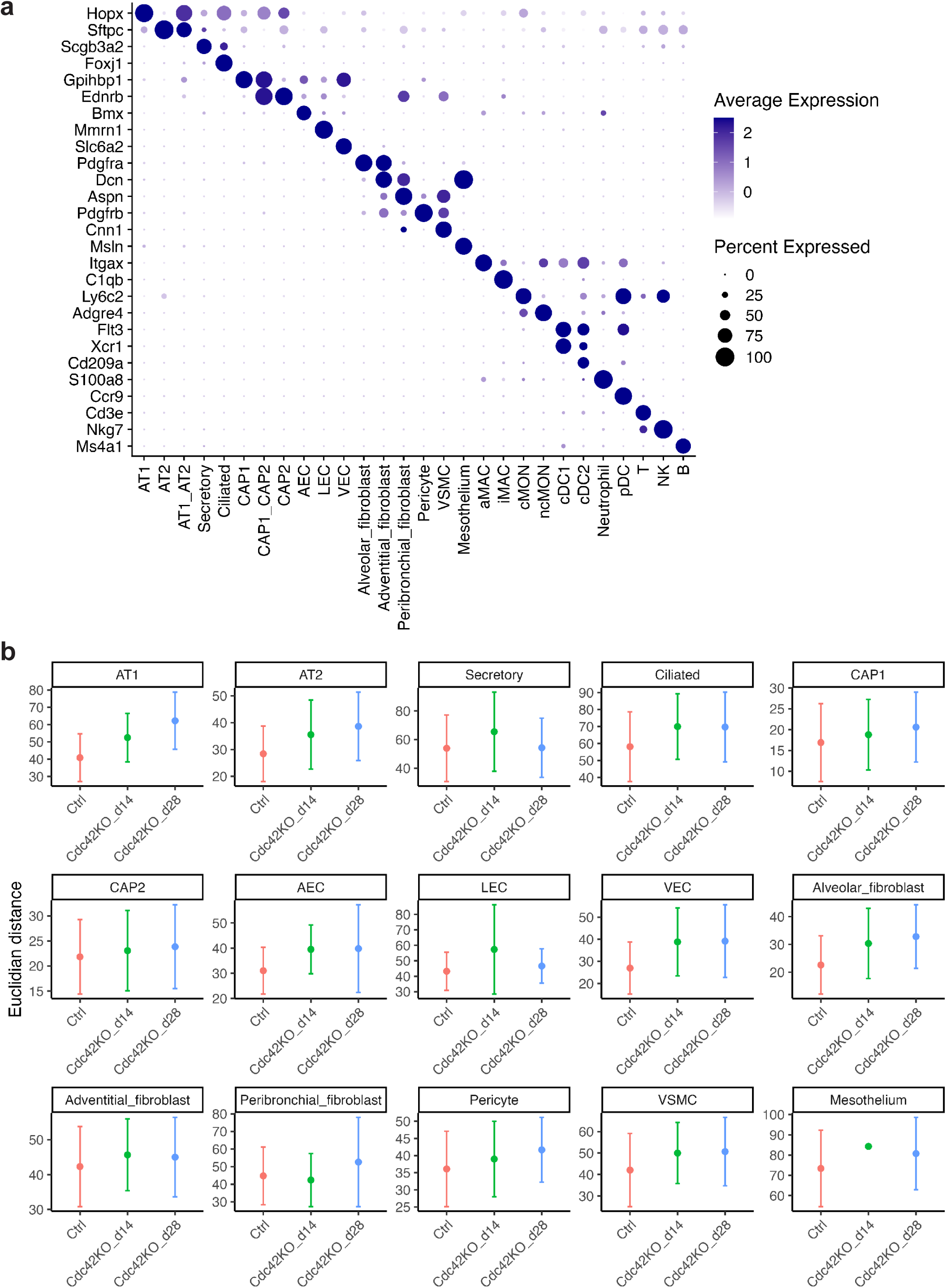
Perturbed mechanosignaling in AT1 cells alters gene transcription in neiboring cells within the alveolus. **a.** A dot plot showing marker gene expression in each cell type from the *Cdc42*^AT1-KO^ scRNA seq dataset. A schematic of experiment is shown in Fig. 3a and a UMAP is shown in Fig. 3b. **b.** Euclidian distance indicating cellular transcriptional similarity across treatments in indicated cell types.

**Extended Data Figure 6:**
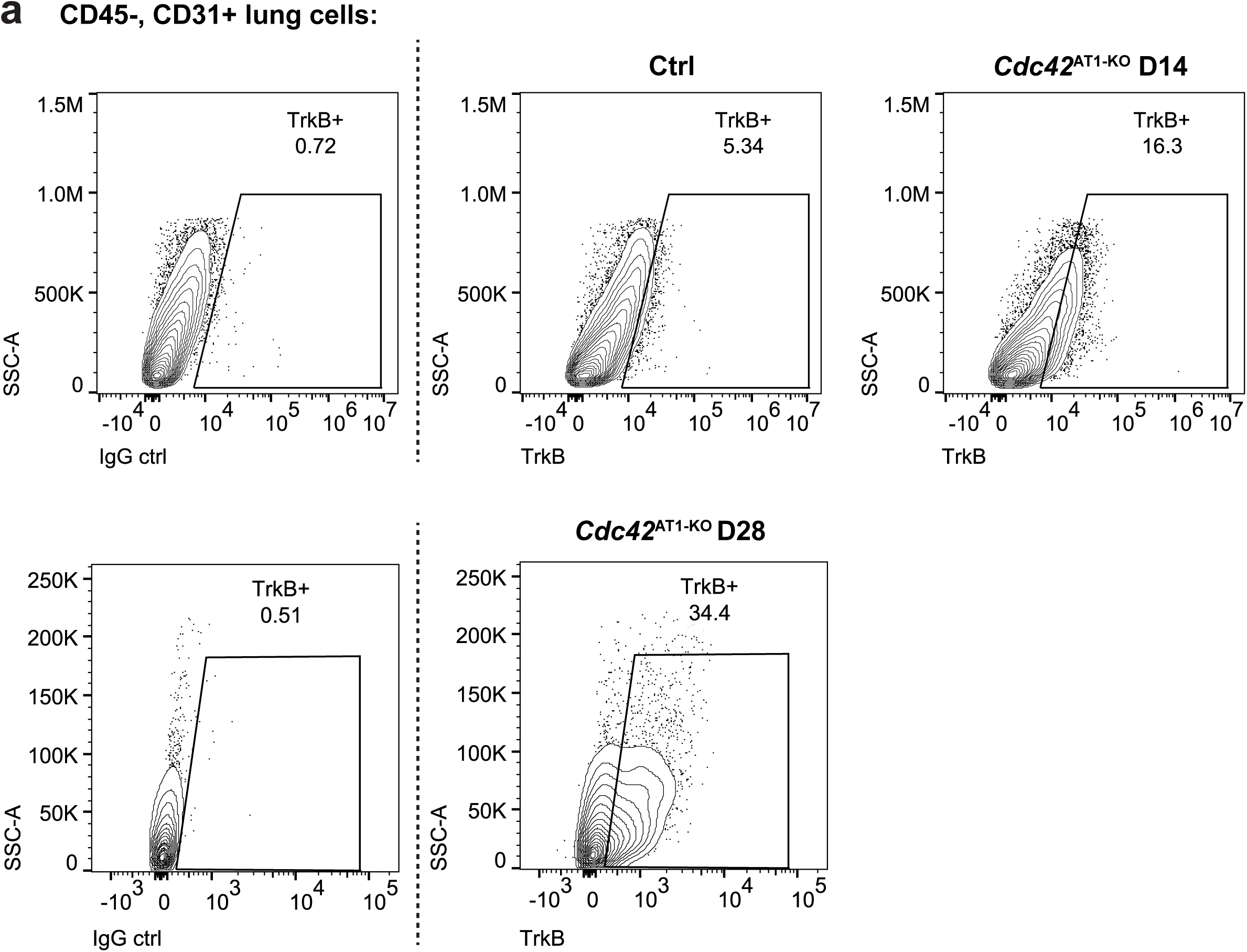
Perturbed mechanosignaling in AT1 cells induces the emergence of TrkB^+^ CAP state. **a.** Representative gating strategies of flow cytometry detecting TrkB^+^ ECs. A quantification of TrkB ^+^ ECs in each condition is shown in Fig. 3h.

**Extended Data Figure 7:**
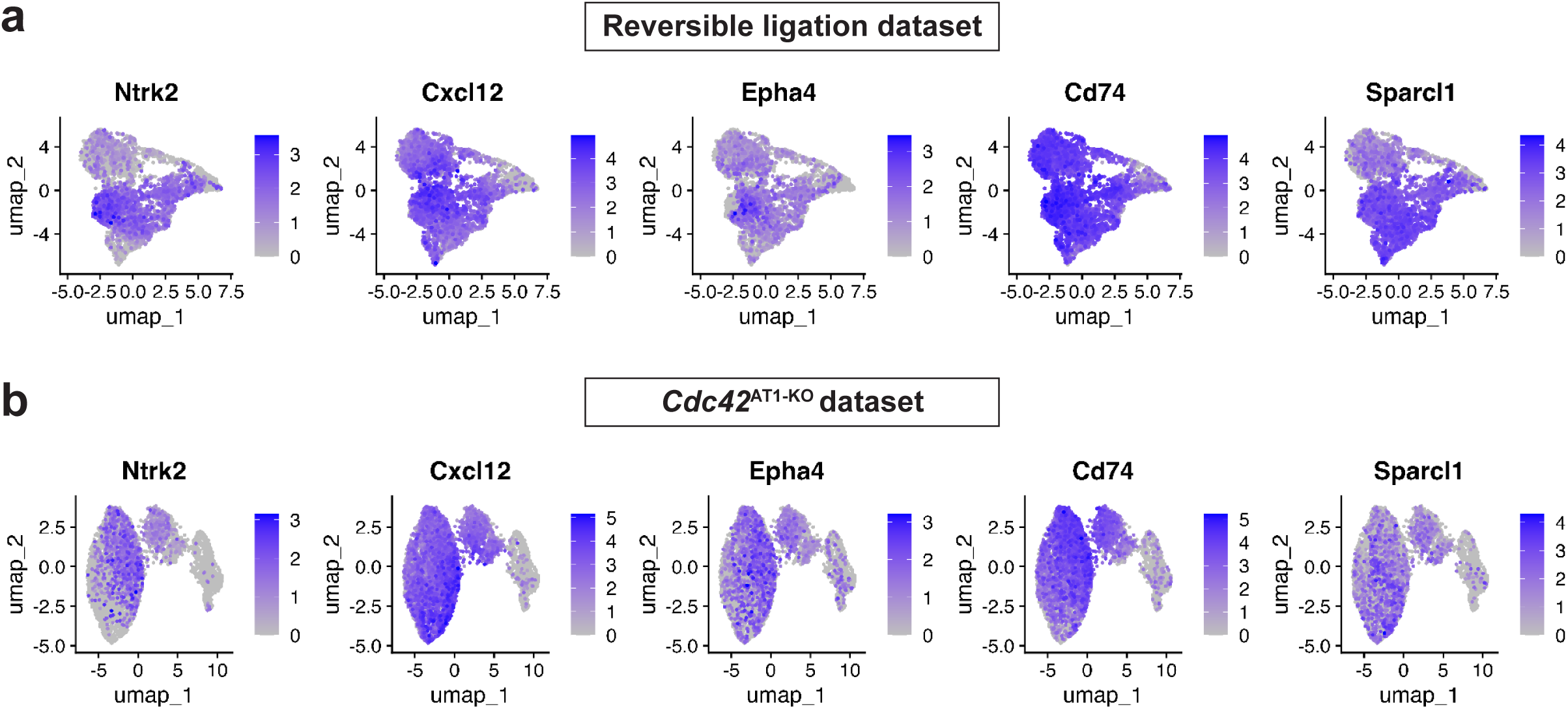
Expression of signature genes in miCAPs. **a-b.** Feature plots showing expression of miCAP signature genes in CAP subset from (**a**) reversible ligation or (**b**) *Cdc42*^AT1-KO^ scRNA seq dataset.

**Extended Data Figure 8:**
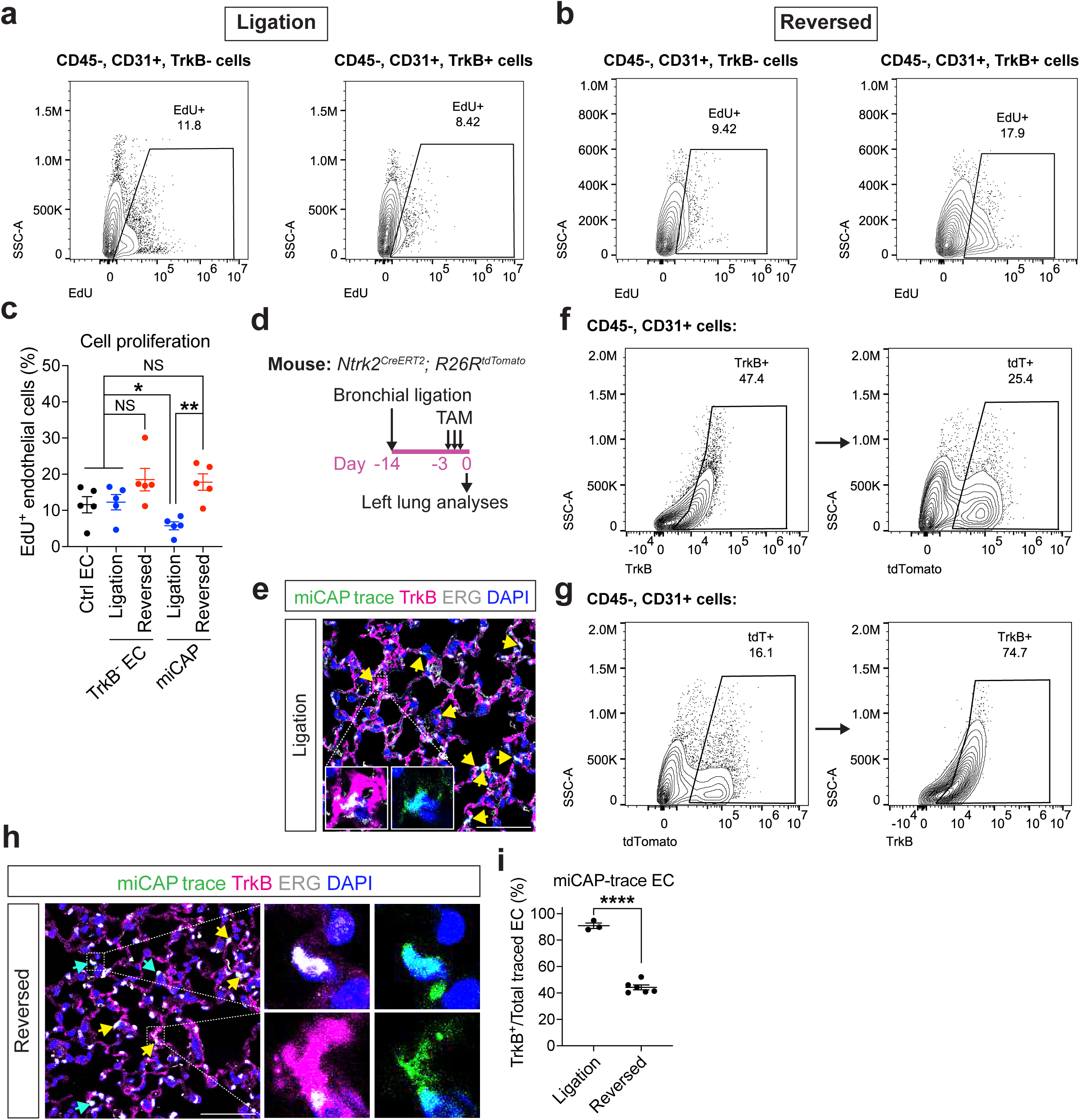
The miCAP state is functionally distinct. **a-b.** Representative gating strategies of flow cytometry detecting EdU^+^ miCAPs. A schematic of experiment is shown in Extended Data Fig. 3a. **c.** Quantification of EdU^+^ miCAPs or TrkB^-^ ECs in each condition (n = 5, mean ± S.E.M.). Asterisks indicate statistical significance (* *p* value < 0.05, ** *p* value < 0.01, unpaired two-tailed Student’s *t* test). NS: no statistical significance. **d.** A schematic of miCAP lineage labelling. *Ntrk2^CreERT2^; R26R^tdTomato^* mice received bronchial ligation surgery, followed by three doses of tamoxifen before euthanasia. Cells and tissues from ligated mouse lungs were used for flow cytometry and immunostaining, respectively. **e.** Representative IHC images of indicated lung tissues stained with RFP (miCAP lineage trace marker), TrkB, and ERG (scale bar = 25 μm). Yellow arrows indicated miCAP lineage^+^ ECs that marked by TrkB expression. **f-g.** Representative flow cytometry gating showing (**f**) efficiency and (**g**) specificity of miCAP lineage trace model. **h.** Representative IHC images of indicated lung tissues stained with RFP (miCAP lineage trace marker), TrkB, and ERG (scale bar = 25 μm). Yellow arrows indicated miCAP lineage^+^ ECs that retained TrkB expression, whereas cyan arrows indicated those that lost TrkB expression. **i.** Quantification showing miCAP-trace ECs that are positive for TrkB expression after ligation or reversed surgery (n = 3 or 6, mean ± S.E.M.). Asterisks indicate statistical significance (**** *p* value < 0.0001, unpaired two-tailed Student’s *t* test).

**Extended Data Figure 9:**
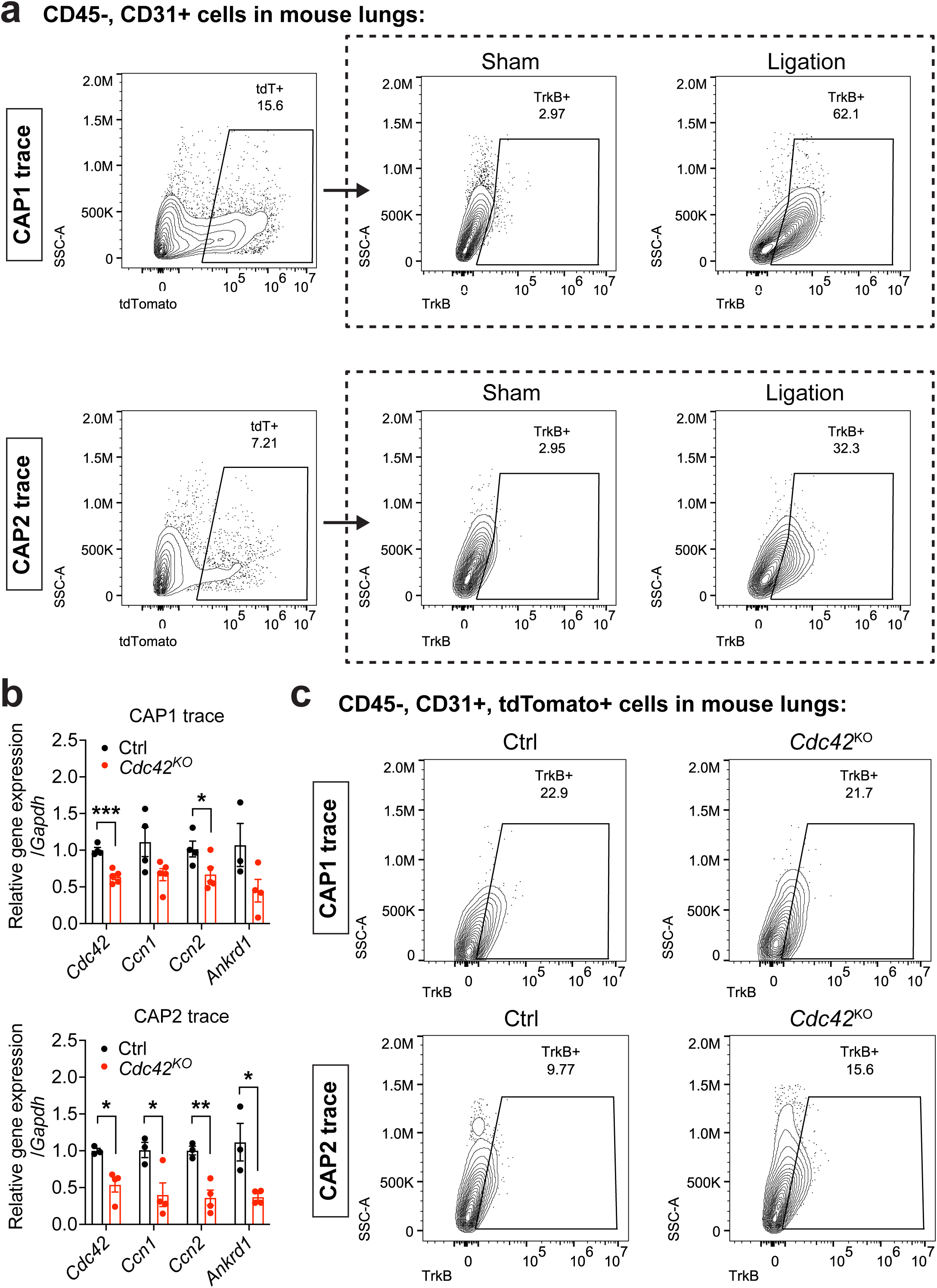
miCAP emergence is independent of intrinsic CAP mechanosignaling. **a.** Representative gating strategies of TrkB^+^ CAP1- or CAP-trace ECs after ligation. A schematic of experiment is shown in Fig. 4j and a quantification is shown in Fig. 4k. **b.** Quantitative RT-PCR showing the downregulation of mechanosignaling sensitive genes (*Ccn1*, *Ccn2*, *Ankrd1*) after *Cdc42* knockout in CAP1 or CAP2s (n = 4 or 5, mean ± SEM). *Ankrd1* expression was not detected in CAP1 and CAP2 after ligation (n = 1 from each group). Asterisks indicate statistical significance (* *p* value < 0.05, ** *p* value < 0.01, *** *p* value < 0.001, unpaired two-tailed Student’s *t* test). **c.** Representative gating strategy of TrkB^+^ CAP1- or CAP2-trace ECs after lineage-specific *Cdc42* deletion. A schematic of experiment is shown in Fig. 4m and a quantification is shown in Fig. 4n.

**Extended Data Figure 10:**
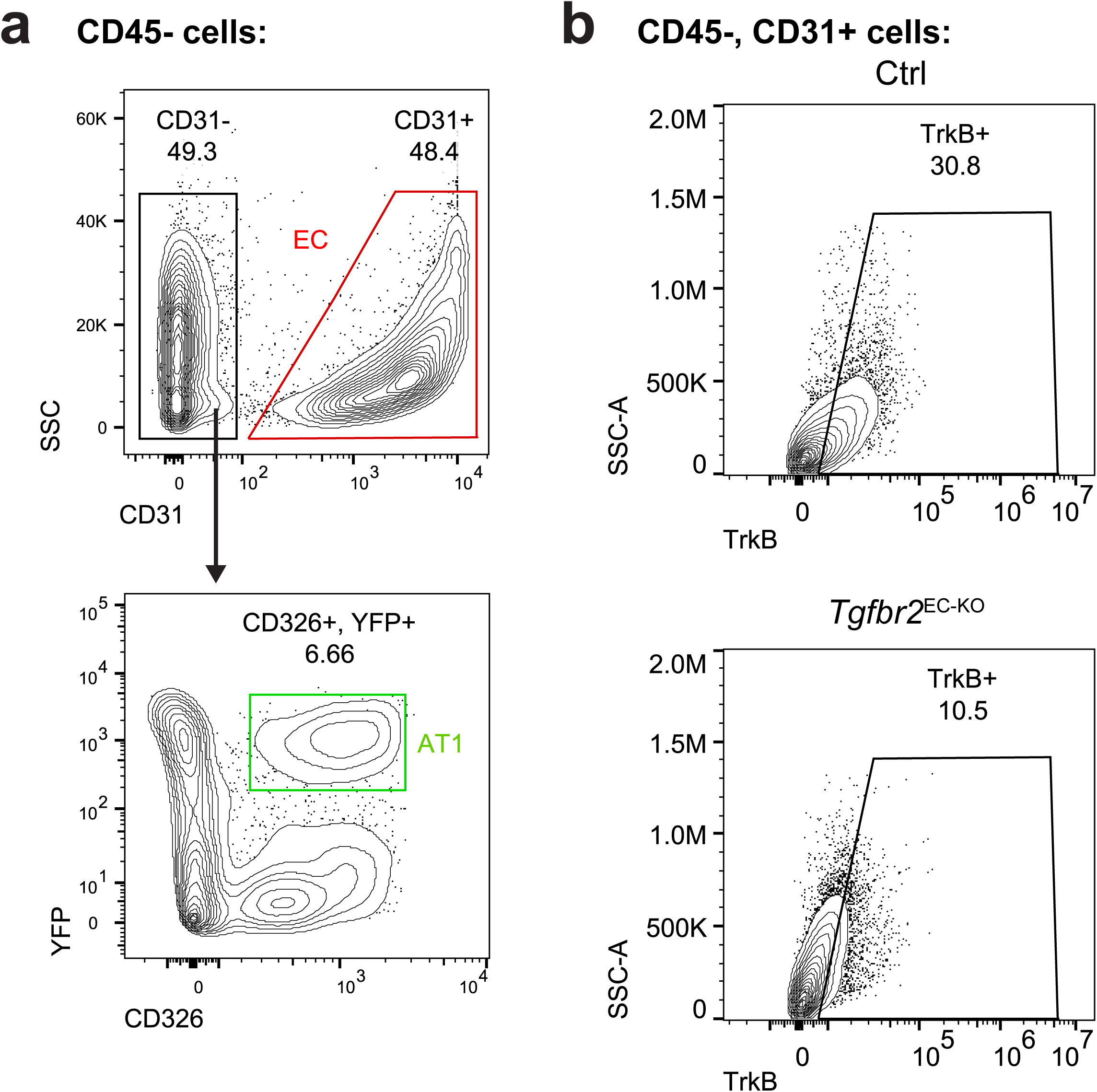
Representative FACS and flow cytometry gating strategies. **a.** Representative gating strategies to isolate lung ECs and AT1 cells by FACS. **b.** Representative flow cytometry gating of TrkB^+^ ECs after ligation in control or *Tgfbr2*^EC-KO^ mouse lungs. A schematic of experiment is shown in Fig. 5d and a quantification is shown in Fig. 5e.

**Extended Data Figure 11:**
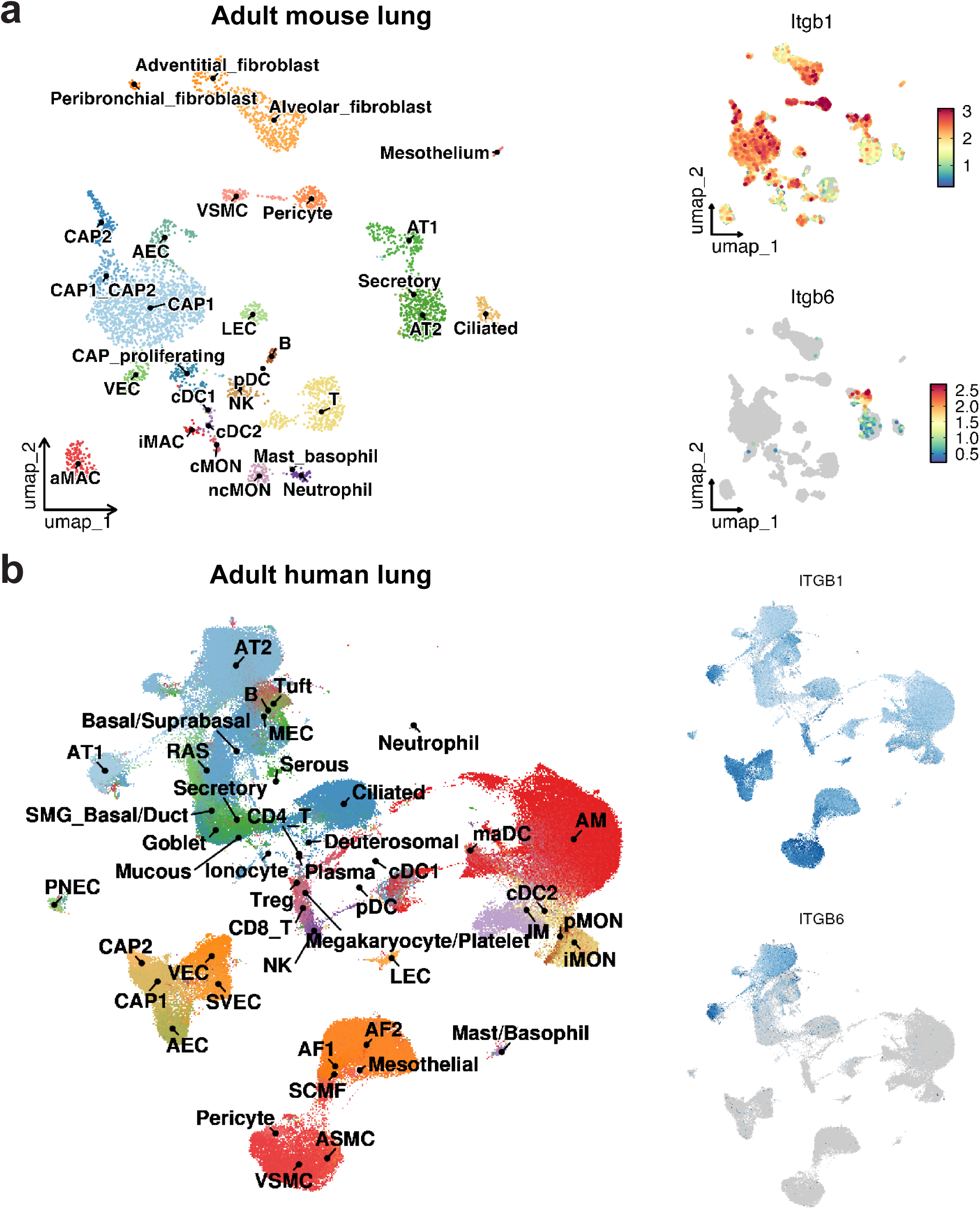
Integrin β6 is specifically expressed in AT1 cells. **a-b.** UMAP plots showing expression of the gene encoding integrin β6 in wildtype (**a**) mouse or (**b**) human lungs. Data in **a.** was also shown in Fig. 2b.

**Extended Data Figure 12:**
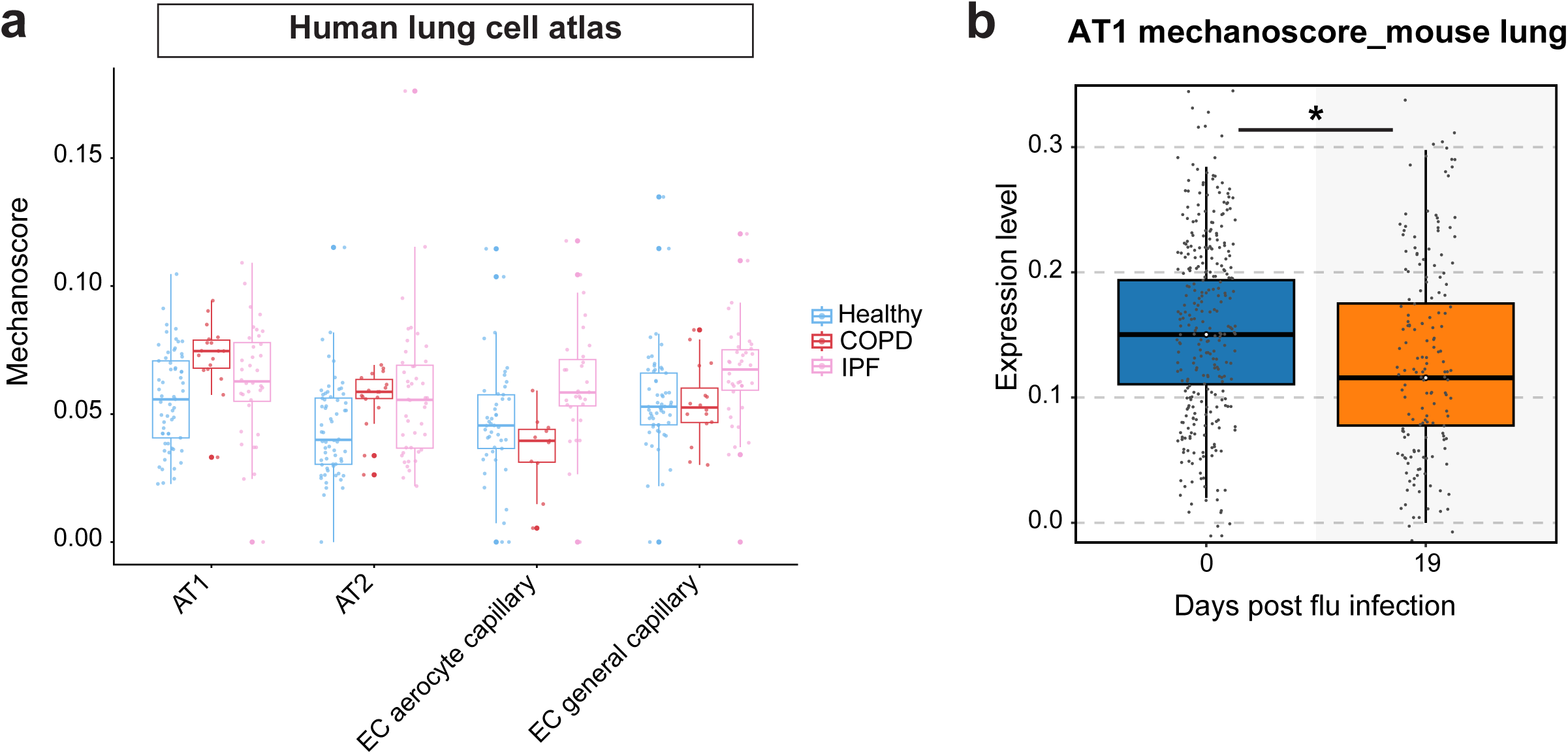
AT1 mechanosignaling is altered in acute lung injury and chronic lung diseases. **a.** Mechanoscore of indicated cell types in the tissue from human healthy, COPD, and IPF lungs (Habermann *et al*. 2020, GSE135893; Adams *et al*. 2020, GSE136831) generated using pseudobulk aggregated expression of YAP/TAZ target genes (median ± S.D.). **b.** Mechanoscore of indicated cell types in control and influenza infected mouse lungs (Niethamer *et al*. 2025, GSE262927). An asterisk indicates statistical significance (* *p* value < 0.05, pairwise Wilcoxon test).

